# Inhibiting Spinal Cord-Specific Hsp90 Isoforms Reveals a Novel Strategy to Improve the Therapeutic Index of Opioid Treatment

**DOI:** 10.1101/2021.04.14.439852

**Authors:** David I. Duron, Christopher S. Campbell, Kerry Chou, Parthasaradhireddy Tanguturi, Paul Bejarano, Katherin A. Gabriel, Jessica L. Bowden, Sanket Mishra, Christopher Brackett, Deborah Barlow, Karen L. Houseknecht, Brian S.J. Blagg, John M. Streicher

**Affiliations:** Department of Pharmacology, College of Medicine, University of Arizona, Tucson AZ USA; Department of Chemistry and Biochemistry, College of Science, University of Notre Dame, Notre Dame IN USA; Department of Biomedical Sciences, College of Osteopathic Medicine, University of New England, Biddeford ME USA; Comprehensive Pain and Addiction Center, University of Arizona, Tucson AZ USA

**Keywords:** Heat shock protein 90, opioid, anti-nociception, reward, constipation, tolerance, therapeutic index, isoforms

## Abstract

Opioid drugs like morphine are the gold standard for the treatment of chronic pain, but are limited by adverse side effects, such as tolerance, constipation, and reward/addiction. In our earlier work, we showed that Heat shock protein 90 (Hsp90) has a crucial role in regulating opioid signaling that differs between brain and spinal cord; Hsp90 inhibition in brain blocks opioid pain relief, while inhibition in the spinal cord enhances it. Building on these findings here, we injected the non-selective Hsp90 inhibitor KU-32 directly into the spinal cord of male and female CD-1 mice, showing that morphine anti-nociceptive potency was boosted by 1.9-3.5 fold in the pain models of tail flick, post-surgical paw incision, and HIV peripheral neuropathy. At the same time, morphine tolerance was reduced from 21 fold to 2.9 fold and established tolerance was rescued, while the potency of constipation and reward (as measured by conditioned place preference) was unchanged. These results demonstrate that spinal Hsp90 inhibition can improve the therapeutic index of morphine. However, we also found that systemic non-selective Hsp90 inhibition resulted in a brain-like effect, blocking opioid pain relief. We thus sought a way to circumvent the effects of brain Hsp90 inhibition by investigating the molecular Hsp90 isoforms active in regulating opioid signaling in both regions. Using selective small molecule inhibitors and CRISPR gene editing, we found that 3 Hsp90 isoforms regulated spinal cord opioid signaling (Hsp90α, Hsp90β, and Grp94) while our previous work showed only Hsp90α was active in brain. We thus hypothesized that a systemically delivered selective inhibitor to Hsp90β or Grp94 could selectively inhibit spinal cord Hsp90 activity, resulting in enhanced opioid pain relief and decreased side effects. We tested this hypothesis using intravenous delivery of KUNB106 (Hsp90β) and KUNG65 (Grp94), showing that both drugs enhanced morphine potency in tail flick and paw incision pain while rescuing anti-nociceptive tolerance. We also found that intravenous KUNA115 (Hsp90α) fully blocked morphine anti-nociception. Together, these results suggest that selective inhibition of spinal cord Hsp90 isoforms is a novel, translationally feasible strategy to improve the therapeutic index of opioids.

## Introduction

Opioid drugs like morphine are the gold standard for the treatment of moderate to severe chronic pain, but are limited by serious side effects, including tolerance, constipation, and reward/addiction liability [1, 2]. These limitations have spurred the search for alternate approaches to improve opioids, such as multifunctional ligands, or opioid agonists biased against the recruitment of βarrestin2 (recently reviewed in [3]). While none of these approaches has solved the opioid issue, they have revealed how manipulating the signal transduction cascades of the opioid receptors holds great promise in improving opioid outcomes (e.g. arrestin bias).

To this end, we’ve engaged in a long-term effort to investigate the role of Heat shock protein 90 (Hsp90) in regulating opioid signaling and anti-nociception. Hsp90 is a ubiquitous and highly expressed chaperone protein with a variety of roles, including nascent protein maturation, signaling kinase activation, and scaffolding signaling complex formation (reviewed in [4, 5]). Hsp90 has mostly been studied in the context of cancer. A few papers have shown that Hsp90 promotes inflammation during inflammatory and neuropathic pain [6–8], and another two papers have shown that Hsp90 could promote opioid dependence and withdrawal [9, 10]. More recently, another paper has linked the Hsp90β isoform to opioid receptor signaling [11]. However, in general, the role of Hsp90 in the pain and opioid systems is mostly unstudied (reviewed in [12]).

In our work, we’ve uncovered a role for Hsp90 in opioid signaling and anti-nociception that differs between the brain and spinal cord. In brain, Hsp90 promotes ERK MAPK signaling via the Hsp90α isoform and the cochaperones Cdc37 and p23; Hsp90 inhibitor treatment in the brain thus reduces ERK MAPK signaling and opioid anti-nociception [13–15]. In contrast, Hsp90 represses opioid anti-nociception and signaling in the spinal cord, so that Hsp90 inhibitor treatment in the spinal cord promotes an ERK-RSK signaling cascade that results in enhanced opioid anti-nociception [16].

This enhanced anti-nociception led us to hypothesize that spinal Hsp90 inhibitor treatment could be used to improve the therapeutic index of opioids and enable a dose-reduction strategy. This is because many side effects are regulated outside the spinal cord, and would presumably not be impacted by spinal inhibition (e.g. reward in ventral tegmental area and striatum [17], constipation in the gut [18]). If anti-nociception were improved but side effects were either improved or not altered, then a lower dose of opioid could be given in combination with Hsp90 inhibitor treatment. This would hypothetically result in improved or maintained anti-nociception with decreased side effects (see [19] for a parallel approach). A novel dose-reduction strategy of this kind could be used to improve opioid therapy in chronic pain patients, decreasing the impact of the negative side effects of opioid therapy.

## Materials and Methods

### Drugs and CRISPR constructs

KU-32, KUNA115, KUNB106, and KUNG65 were synthesized, purified, and characterized as in our previously published work (KU-32 is Compound A4 in [20]; KUNA115 in [21]; KUNB106 in [22]; KUNG65 in [23]). Purity was confirmed by HPLC (>95%) and identity confirmed by HRMS and NMR. Inhibitors were stored under desiccation at −20°C, and stock solutions were prepared in DMSO, and also stored at −20°C. A matched Vehicle control for was included in every experiment; 0.02% DMSO in sterile USP water for the 0.01 nmol intrathecal injections and 10% DMSO, 10% Tween80, and 80% sterile USP saline for 1 mg/kg intravenous and 10 mg/kg oral injections. Morphine sulfate pentahydrate was obtained from the NIDA Drug Supply Program, stored at room temperature, and working solutions were made fresh prior to every experiment in sterile USP saline. USP saline injected controls were used for the reward and constipation assays below.

All-in-one CRISPR DNA constructs expressing Cas9 and a gRNA targeting Hsp90α (MCP229411-CG01-3-B), Hsp90β (MCP227368-CG12-3-B), and Grp94 (MCP230394-CG12-3-B) were obtained from Genecopoeia (Rockville, MD). The DNA was amplified using standard molecular biology approaches, and complexed with TurboFect *in vivo* transfection reagent (Thermo Fisher, Waltham, MA) as described in our previous work [14] and by the manufacturer’s protocol. The complexed DNA was injected into the mice by the intrathecal route (2 μg DNA in 5 μL) daily from days 1-3, with behavioral testing performed on day 10.

### Mice

Male and female CD-1 mice in age-matched cohorts from 5–8 weeks of age were used for all behavioral experiments and were obtained from Charles River Laboratories (Wilmington, MA). CD-1 (a.k.a. ICR) mice are commonly used in opioid research as a line with a strong response to opioid drugs (*e.g.* [24], and our own previous Hsp90 research [13–16]). Mice were recovered for a minimum of 5 days after shipment before being used in experiments. Mice were housed no more than 5 mice per cage and kept in an AAALAC-accredited vivarium at the University of Arizona under temperature control and 12-h light/dark cycles. All mice were provided with standard lab chow and water available *ad libitum*. The animals were monitored daily, including after surgical procedures, by trained veterinary staff. All experiments performed were in accordance with IACUC-approved protocols at the University of Arizona and by the guidelines of the NIH Guide for the Care and Use of Laboratory Animals. We also adhered to the guidelines of ARRIVE; no adverse events were noted for any of the animals.

### Behavioral experiments

All animals were randomized to treatment groups by random assignment of mice in one cohort to cages, followed by random block assignment of cages to treatment group. Group sizes were based on previous published work from our lab using these assays [14, 16, 25, 26]. The mice were not habituated to handling. Prior to any behavioral experiment or testing, animals were brought to the testing room in their home cages for at least 1 h for acclimation. Testing always occurred within the same approximate time of day between experiments during the animal light (inactive) cycle, and environmental factors (noise, personnel, and scents) were minimized. All testing apparatus (cylinders, grid boxes, etc.) were cleaned between uses using 70% ethanol and allowed to dry. The experimenter was blinded to treatment group by another laboratory member delivering coded drug vials, which were then decoded after collection of all data. Naïve mice were used for every experiment, including each dose.

### Paw incision and mechanical allodynia

Mechanical thresholds were determined prior to surgery using calibrated von Frey filaments (Ugo Basile, Varese, Italy) with the up-down method and four measurements after the first response per mouse as in [27] and our previously published work (e.g. [28]). The mice were housed in a homemade apparatus with Plexiglas walls and ceiling and a wire mesh floor (3-inch wide 4-inch long 3-inch high with 0.25-inch wire mesh). The surgery was then performed by anesthesia with ~2% isoflurane in standard air, preparation of the left plantar hind paw with iodine and 70% ethanol, and a 5-mm incision made through the skin and fascia with a no. 11 scalpel. The muscle was elevated with curved forceps leaving the origin and insertion intact, and the muscle was split lengthwise using the scalpel. The wound was then closed with 5-0 polyglycolic acid sutures. Mice were then injected with inhibitor or Vehicle control and left to recover for 24 h. Our intrathecal (i.t.) injection protocol is reported in [13]; briefly, the injection was made in awake and restrained animals with a 10 μL Hamilton syringe and 30 g needle (5-7 μL volume) between the L5-L6 vertebrae at a 45° angle, with placement validated by tail twitch. Since the injection was made through skin with no incision or other surgical intervention, repeated injection protocols were performed the same way. The next day, the mechanical threshold was again determined as described above. Mice were then injected with 0.32-5.6 mg/kg morphine by the subcutaneous (s.c.) route, and mechanical thresholds were determined over a 3-hour time course. No animals were excluded from these studies.

### Tail-flick assay

Pre-injection tail-flick baselines were determined in a 52°C tail-flick assay with a 10-s cutoff time (method also reported in [13]). The mice were then injected with inhibitor or Vehicle control with a 24-hour treatment time. 24-hours post-injection baselines were determined. The mice were then injected s.c. with 1-10 mg/kg of morphine, and tail-flick latencies were determined over a 2-hour time course. For tolerance studies, baseline tail flick latencies were taken, and mice were then injected with inhibitor or Vehicle control with a 24-hour treatment time. 24 hours later mice were baselined again and then injected with 10 mg/kg s.c. morphine with one tail flick latency measured at 30 minutes post morphine. Mice were injected again with inhibitor or Vehicle and the process was repeated for an additional 7 days with twice daily morphine injection, and tail flick response measured after the morning injection. No animals were excluded from these studies.

### HIV peripheral neuropathy

Mechanical threshold baselines were measured prior to any treatment on the left hind paw using von Frey filaments. HIV peripheral neuropathy was induced by intrathecal injection of gp120 IIIb protein (SPEED BioSystems, Gaithersburg, MD, Cat# YCP1549, 15 ng/μl in 0.1 M PBS and 0.1% BSA, 7-μl volume) using our previously established protocol [13] on days 1, 3, and 5. On day 20 a second mechanical threshold baseline was measured on the left hind paw using von Frey filaments and then KU-32 or Vehicle was injected i.t with a 24-h treatment time. A third mechanical threshold was then measured on day 21 and morphine (0.32-10 mg/kg s.c.) was then injected, and mechanical thresholds were measured over a time course on the left hind paw. No animals were excluded from these studies.

### Conditioned place preference

Conditioned place preference training, baseline runs, and post-training runs were all performed in Spatial Place Preference LE 896/898 rigs (Harvard Apparatus, Holliston, MA). Rigs were designed to consist of two chambers with one connecting chamber. Of the two conditioned chambers, one consisted of black and grey dotted walls with a textured floor. The other chamber consisted of black and grey striped walls with smooth floor. Chamber floors connected to a pressure sensor which transferred ongoing data to a computer running PPC WIN 2.0 software (Harvard Apparatus). Prior to preference training baselines were taken on day 0. Mice were placed in CPP chambers and allowed to roam freely for 15 minutes at ~7am. Chambers were cleaned thoroughly with VersaClean and allowed to dry in-between mice. Mice were then injected with i.t. KU-32 or Vehicle with a 24hour treatment time. On day 1 mice were injected with i.t. KU-32 or Vehicle again and allowed to recover for 30 minutes. Mice were then injected s.c. with saline or morphine (3.2, 5.6, or 10mg/kg) at ~7am and placed in either stripe or dotted chambers. Half of each group paired morphine with the striped chamber and the other half to the dotted chamber in an unbiased design. At ~12pm mice were then given a second injection of either saline or morphine which was paired to the opposite chamber. This training process was repeated for 4 days total with morning and noon pairings alternating each day. On day 5 mice were placed in CPP chambers and allowed to roam freely for 15 minutes at ~7am. Raw data in the form of seconds and percentage spent in each chamber was exported from PPC WIN 2.0 as an excel file and transferred to GraphPad Prism 9.3 (San Diego, CA) for further analysis.

### Opioid induced constipation

Prior to the experiment mice were injected with either KU-32 or Vehicle i.t. and allowed to recover for 24 hours. Morphine (1, 3.2, or 10mg/kg s.c.) or saline was injected and followed by a 6-hour fecal production time course. During this time course the mice were housed in the von Frey boxes used to collect the paw incision and HIV neuropathy data above, which have a grate above a collection plate. The feces were counted and weighed in 1-hour bins and used to construct a cumulative plot. Morphine treated groups were normalized to saline groups and represented as a percentage at each timepoint.

### Respiratory depression

Respiratory activity and subsequent morphine-induced depression was measured using whole body plethysmography in awake and freely moving mice using chambers from Data Sciences International (St. Paul, MN). Chambers were maintained at room temperature, with composition of the atmosphere set by mass flow controllers. Mice were injected with inhibitors or Vehicle control, followed by 24 hr treatment time, as for the above assays. The mice were then placed in the chambers for a 30 minute acclimation and baseline period, with respiratory measurements recorded for the last 7 minutes of the period. All mice were then treated with 7.5 mg/kg morphine i.v., immediately placed back in the chambers, and respiratory activity recorded for an additional hour.

### In vitro *ADME assays*

#### LogD Determination

Distribution coefficient (LogD) was determined by the method of Wilson et al. [29]. Briefly, 5 μL volumes of test compounds diluted in DMSO were added to a mixture of equal volumes of 50 mM phosphate buffer and 1-octanol. The compounds were assayed using a 50 nM concentration in order to limit compound precipitation and to ensure that the assay values were maintained within the dynamic range of the LC-MS/MS instrumentation. Samples were vortex mixed at 800 rpm for 24 h. Subsequent to centrifugation at 14000 rpm for 30 min, 1 μL of each layer was analyzed by LC-MS/MS. LogD was calculated using the peak areas obtained from each layer.

#### Aqueous Solubility

Aqueous solubility was determined using a miniaturized shake flask approach, under conditions of pH 6.8 and analyte concentration of 1.0 mM by the method of Zhou et al. [30]. Aqueous solutions of analyte were incubated at room temperature in the chamber of a Whatman (Piscataway, NJ) Mini-UniPrep syringeless filter for 24 h while shaking gently (600 rpm). Subsequent to incubation, filter plungers were pushed down to the bottom of the syringeless filter chamber assemblies, allowing filtrate to enter the plunger compartment. Following an additional 30 min incubation at room temperature, filtrates were diluted with 50:50 acetonitrile/water + 0.1% formic acid and analyzed by LC-MS/MS. Analyte concentrations were determined by the interpolation of peak area ratio from a calibration curve formed by matrix spiked with authentic reference material.

#### *In Vitro* Mouse Liver Microsomal Stability

Metabolic stability of lead compounds was assessed *in vitro* by the method of Di et al. [31]. Briefly, mouse liver microsomes (Corning Life Sciences, Woburn, MA) were isolated from CD-1 mice (male mice, 8-10 weeks of age). Assays were conducted using 0.123 mg/mL protein concentration (total protein concentration in the microsomal solution) and 1.0 μM drug concentration under incubation conditions of 37 °C. Metabolic stability was determined following 0, 5, 15, 30, and 60 min of incubation time. The samples were analyzed by reversed phase LC using a triple quadrupole mass spectrometer. Compound specific transitions of parent ion to product ion were monitored and percent remaining calculated based on peak area of 5-60 min time points (relative to time zero). Half-life calculations were determined using the formula t½ = −ln(2)/k, where k (min-1) is the turnover rate constant (the slope) estimated from a log-linear regression of the percentage compound remaining versus time.

#### *In Vitro* Human Liver Microsomal Stability

Metabolic stability of the compounds in human liver microsomes was determined by the method described above using pooled human liver microsome preparations from 20 male donors (Corning Life Sciences, Woburn, MA). Assays were conducted as described above and the *in vitro* half-life of the compounds was calculated.

#### LC-MS/MS Analysis

LC-MS/MS analysis was conducted using an Agilent (Santa Clara, CA) 6460 triple quadrupole mass spectrometer coupled with an Agilent liquid chromatography (LC) system. The LC system consists of a binary pump, degasser, column heater, and autosampler. Chromatographic separation was performed on a Waters Atlantis T3 3μm 3.0 x 50 mm analytical column using a ballistic gradient of mobile phase consisting of 0.1% formic acid in water (A) and 0.1% formic acid in acetonitrile (B) at a flow rate of 0.75 mL/min. The mobile phase was heated to a temperature of 45 °C.

### In vivo *pharmacokinetic study*

Mice were dosed with 10 mg/kg KUNG65 by the oral route, with groups over a 2 hr time course. The mice from each time point were sacrificed, and whole blood collected into EDTA tubes from a cardiac puncture. This was followed by saline perfusion to clear the vasculature, and the spinal cords were dissected and snap frozen in liquid nitrogen. The whole blood was separated and the plasma stored in EDTA tubes; all samples were stored at −80°C prior to analysis. Compound exposure in plasma subsequent to *in vivo* dosing was determined by precipitation of protein with acetonitrile. Following centrifugation, the supernatant was injected for LC-MS/MS analysis, and plasma concentration determined via interpolation of peak areas from a standard curve prepared in plasma.

### Competition radioligand binding

KUNB106 binding to the opioid receptors was performed substantially as in our previous work [32–36]. Human mu (#ES-542-C), delta (#RBHODM-K), and kappa (#ES-541-C) opioid-expressing CHO cell lines from PerkinElmer (Waltham, MA) were used. The cells were grown in in 1:1 DMEM/F12 medium with 10% heat-inactivated fetal bovine serum, 1X penicillin-streptomycin supplement, and 500 μg/mL G418 selection antibiotic. Cell pellets for experiments were collected using 5 mM EDTA in PBS and membrane protein extracted as described in our cited work. For the binding, 25-30 μg of membrane protein was combined with a fixed concentration (0.57-5.18 nM) of ^3^H-diprenorphine (PerkinElmer) and concentration curves of KUNB106 or positive control (naloxone for mu and delta, U50,488 for kappa) in a 200 μL volume. The reactions were incubated for 1 hr at room temp, then collected onto 96 well format GF/B filter plates using a Brandel Cell Harvester (Gaithersburg, MD). The radioactivity was read using a 96 well format MicroBeta2 scintillation counter (PerkinElmer). The resulting data was normalized to binding in the presence of Vehicle (100%) or 10 μM naloxone/U50,488 (0%), and used to calculate the K_I_ based on the previously established K_D_ of ^3^H-diprenorphine in each cell line using GraphPad Prism 9.3 using a 1-site fit model.

### Statistical analysis

All data were reported as the mean ± SEM and normalized where appropriate as described above. The behavioral data from each individual dose was reported raw without maximum possible effect (MPE) or other normalization. Data for dose/response curves (except CPP) was normalized to %MPE using peak effect, except for constipation, which used Area Under the Curve [MPE = (Response-Baseline) / (Threshold – Baseline) * 100]. The CPP dose/response curve was reported as the % Difference Score [Diff Score = % in Paired - % in Unpaired]. Technical replicates and further details are described in the Figure Legends. Potency (A_50_) values were calculated by linear regression using our previously reported method [13], with further details in the Figure Legends, and were reported with 95% confidence intervals. Statistical comparisons of individual dose/response curve time courses were performed using Repeated Measures 2-Way ANOVA with Sidak’s (tail flick, paw incision, HIV neuropathy, tolerance rescue, constipation, respiratory depression) or Tukey’s (tolerance, CPP) *post hoc* tests. The Geisser-Greenhouse correction was used to account for a potential lack of sphericity of the data, permitting valid Repeated Measures ANOVA. ANOVA *post hoc* tests were only performed when ANOVA F values indicated a significant difference, and there was homogeneity of variance (permitting parametric analysis). In all cases, significance was defined as p < 0.05. The group sizes reported represent independent individual mice tested in each assay. All graphing and statistical analyses were performed using GraphPad Prism 9.3. Approximately equal numbers of male and female mice were used for each experiment. Comparison by 2 Way ANOVA using sex as a variable revealed no sex differences in this study, so all male and female mice were combined.

## Results

### Spinal Hsp90 inhibition enhances morphine anti-nociception in multiple pain models

Based on our earlier work [16] we hypothesized that spinal cord Hsp90 inhibition would consistently enhance morphine anti-nociception over a full dose range in different acute and chronic pain models. We tested this using male and female CD-1 mice injected with 0.01 nmol of the non-isoform-selective Hsp90 inhibitor KU-32 or Vehicle control by the i.t. route with a default treatment time of 24 hours (time point based on our earlier work [13–16]). After the 24 hour treatment time, the mice were treated with a dose range of morphine and antinociception was measured.

We first tested acute thermal nociception in uninjured mice using the well-established tail flick model. KU-32 treatment caused a consistent and significant elevation in morphine anti-nociception over the dose range of 1-5.6 mg/kg (**Figure 1A-C**). The response was not different at 10 mg/kg; however, 10 mg/kg is a maximal dose in this assay, and responses were not recorded past the 10 second threshold (**Figure 1D**). Dose/response analysis revealed a potency of 3.0 (2.5 – 3.7; 95% confidence intervals reported in parentheses) mg/kg in Vehicle treated mice and 1.6 (1.3 – 2.0) mg/kg in KU-32 treated mice, representing a 1.9 fold shift improvement in potency (**Figure 1E**).

**Figure 1:**
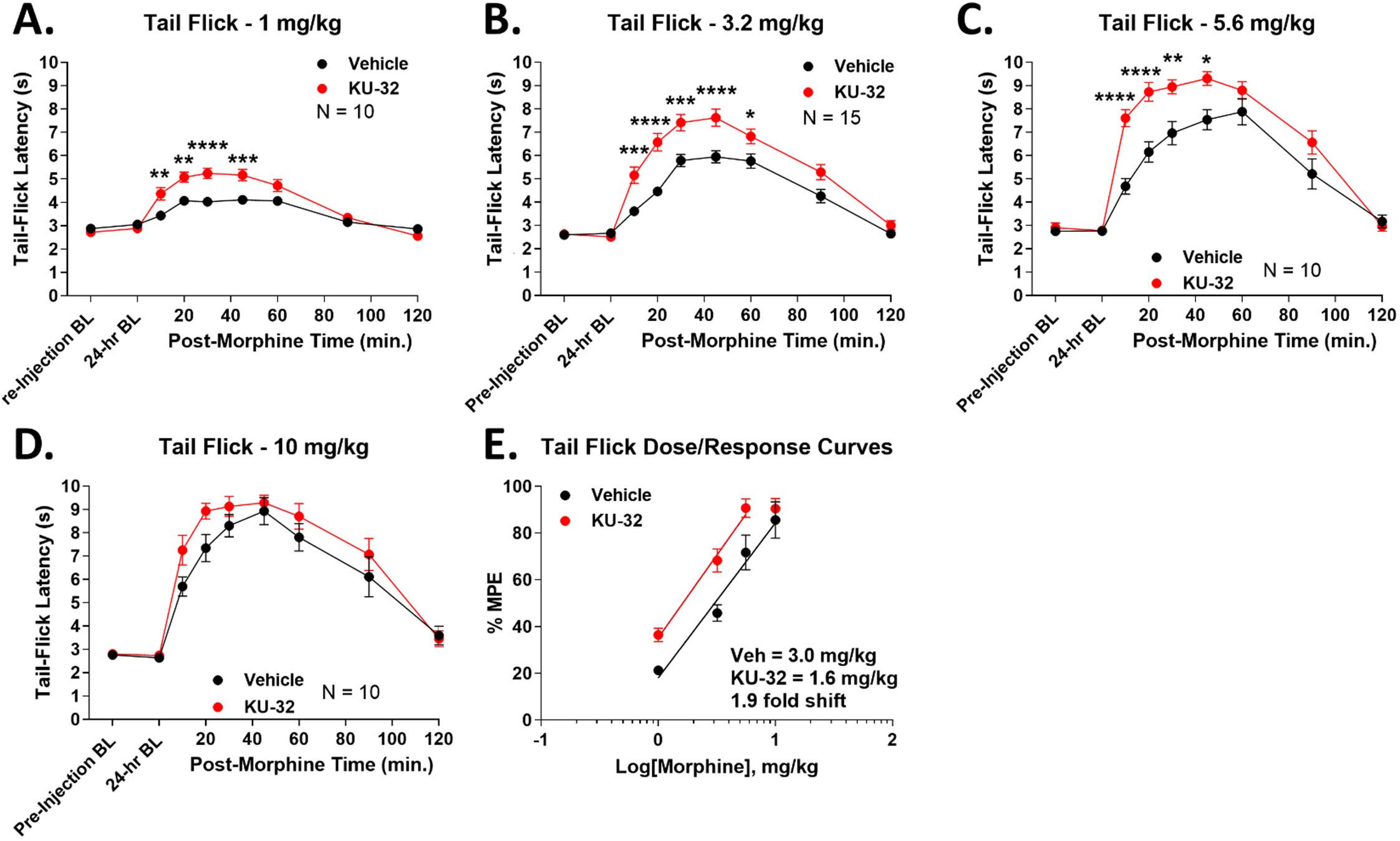
Spinal Hsp90 inhibition increases morphine potency in acute heat-induced tail flick. Male and female CD-1 mice were injected i.t. with 0.01 nmol KU-32 or Vehicle control, 24 hours, then 1-10 mg/kg morphine, s.c.. Tail flick responses were recorded at 52°C with a 10 sec cutoff. Data reported as the mean ± SEM, with sample sizes of mice/group noted in each graph. 2-3 technical replicates were performed per dose. *, **, ***, **** = p < 0.05, 0.01, 0.001, 0.0001 vs. same time point Vehicle group by RM 2 Way ANOVA with Sidak’s *post hoc* test. **A-D)** Individual dose curves shown as noted. Data not normalized. **E)** Dose/response analysis performed for individual curves, normalized as %MPE (baseline vs. 10 sec cutoff). A_50_: Vehicle = 3.0 (2.5 – 3.7) mg/kg; KU-32 = 1.6 (1.3 – 2.0) mg/kg; 1.9 fold increase in potency.

This result was encouraging; however, tail flick is a spinal reflex in uninjured animals, and may not translate to clinical pain conditions. We thus used the post-surgical paw incision assay to model post-surgical pain, a common opioid indication [37]. Much as in heat-induced tail flick, KU-32 caused a significant and consistent elevation in morphine anti-nociception over the 0.32 – 5.6 mg/kg dose range (**Figure 2A-E**). Dose/response analysis showed an A_50_ potency value of 2.4 (2.0 – 2.8) mg/kg for Vehicle animals and 0.85 (0.64 – 1.1) mg/kg for KU-32 animals, representing a 2.8 fold improvement in potency (**Figure 2F**).

**Figure 2:**
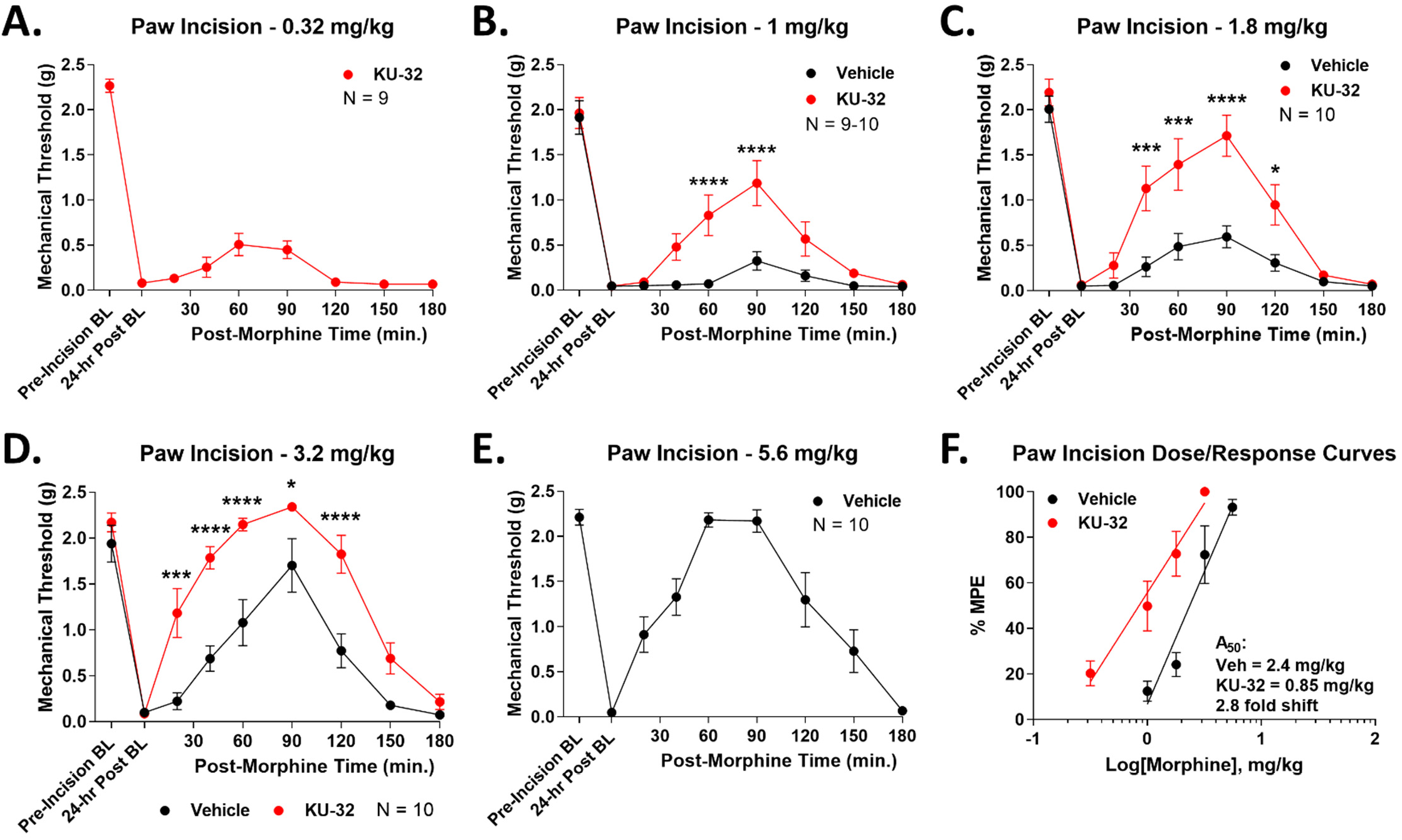
Spinal Hsp90 inhibition increases morphine potency in acute post-surgical paw incision pain. Male and female CD-1 mice had the paw incision surgery performed, then were injected i.t. with 0.01 nmol KU-32 or Vehicle control, 24 hours, then 0.32-5.6 mg/kg morphine, s.c.. Paw withdrawal responses recorded using von Frey filaments, including pre- and post-surgical baselines, validating the pain state. Data reported as the mean ± SEM, with sample sizes of mice/group noted in each graph. 2 technical replicates were performed per dose. *, ***, **** = p < 0.05, 0.001, 0.0001 vs. same time point Vehicle group by RM 2 Way ANOVA with Sidak’s *post hoc* test. **A-E)** Individual dose curves shown as noted. Data not normalized. Vehicle at 0.32 mg/kg and KU-32 at 5.6 mg/kg were not measured since these would be too low (Vehicle) or hit the assay threshold (KU-32), meaning they would not fit the linear dose/response model used. **F)** Dose/response analysis performed for individual curves, normalized as %MPE (baseline vs. 2.34 g cutoff). A_50_: Vehicle = 2.4 (2.0 – 2.8) mg/kg; KU-32 = 0.85 (0.64 – 1.1) mg/kg; 2.8 fold increase in potency.

These results are again promising, but both pain states are acute and short in duration, and do not represent chronic pain, which is the most difficult to treat in the clinic. We thus used the HIV peripheral neuropathy model induced by gp120 protein injection, which is a sustained and long-lasting neuropathic pain model [38]. After 3 sustained weeks of chronic pain, the mice were treated with KU-32 or Vehicle control and tested as above. Again, KU-32 treatment caused a sustained and significant increase in morphine antinociception over the 0.32-10 mg/kg dose range (**Figure 3A-E**). Dose/response analysis showed a potency of 4.2 (3.7 – 4.8) mg/kg for Vehicle treatment and 1.2 (0.87 – 1.5) mg/kg for KU-32 treatment, an improvement of 3.5 fold (**Figure 3F**). Not only did KU-32 improve opioid anti-nociception in this chronic pain model, it had the largest fold-shift improvement of the 3 models tested. Importantly, KU-32 treatment did not cause baseline differences in any pain state measured.

**Figure 3:**
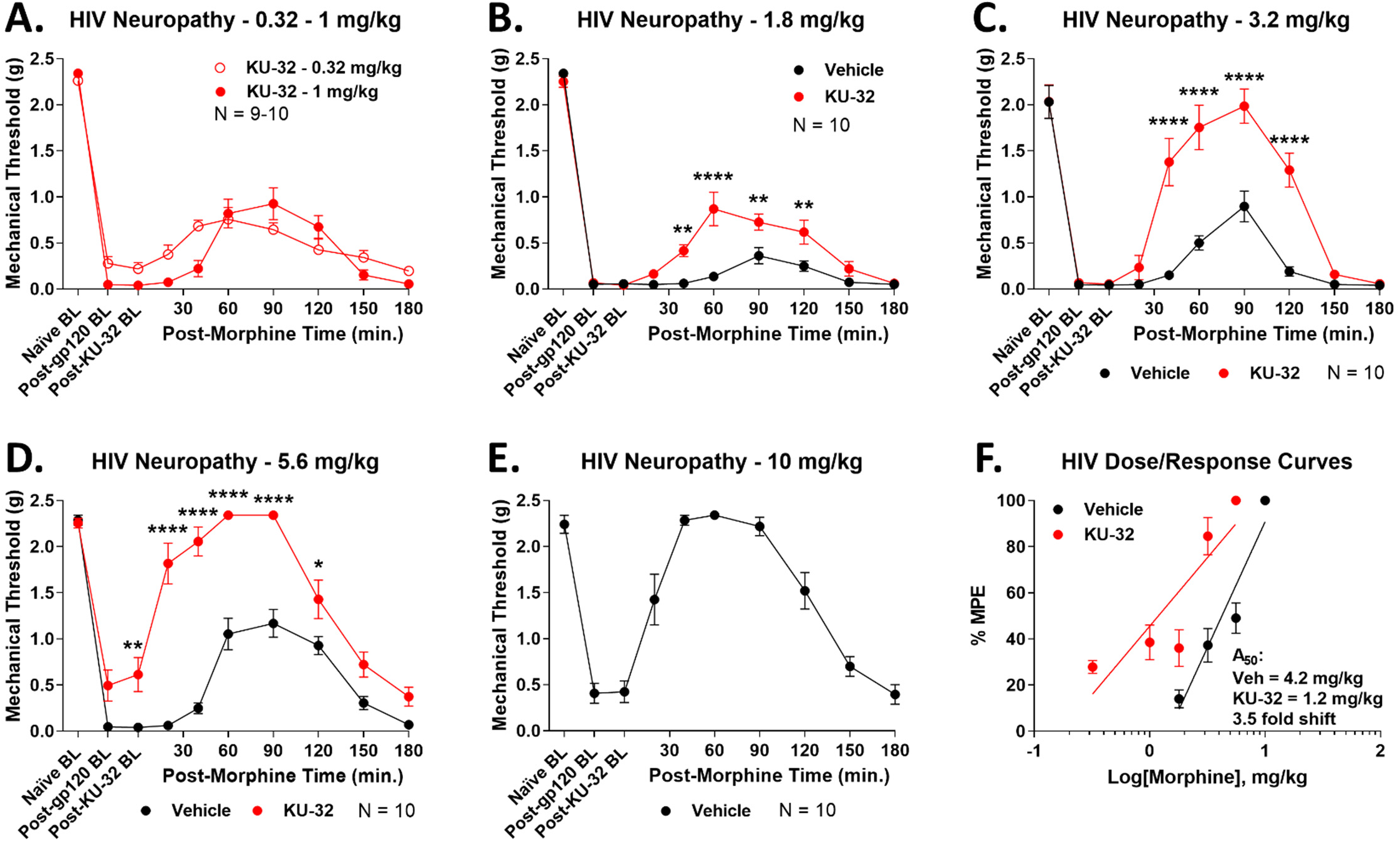
Spinal Hsp90 inhibition increases morphine potency in chronic HIV neuropathy pain. Male and female CD-1 mice were injected i.t. with gp120 protein on days 1, 3, and 5 (see Methods). On day 20, mice were injected i.t. with 0.01 nmol KU-32 or Vehicle control, 24 hours, then 0.32-10 mg/kg morphine, s.c.. Paw withdrawal responses recorded using von Frey filaments, including pre- and post-treatment baselines, validating the pain state. Data reported as the mean ± SEM, with sample sizes of mice/group noted in each graph. 2 technical replicates were performed per dose. *, **, **** = p < 0.05, 0.01, 0.0001 vs. same time point Vehicle group by RM 2 Way ANOVA with Sidak’s *post hoc* test. **A-E)** Individual dose curves shown as noted. Data not normalized. Vehicle at 0.32-1 mg/kg and KU-32 at 10 mg/kg were not measured since these would be too low (Vehicle) or hit the assay threshold (KU-32), meaning they would not fit the linear dose/response model used. **F)** Dose/response analysis performed for individual curves, normalized as %MPE (baseline vs. 2.34 g cutoff). A_50_: Vehicle = 4.2 (3.7 – 4.8) mg/kg; KU-32 = 1.2 (0.87 – 1.5) mg/kg; 3.5 fold increase in potency.

### Spinal Hsp90 inhibition reduces morphine tolerance and rescues established tolerance

The results above support our hypothesis that spinal Hsp90 inhibition could make opioids more potent. However, at the same time, if side effect potencies are enhanced, then the improvement does not result in an improved therapeutic index or improved opioid therapy. We thus tested the potency of morphine side effects with spinal Hsp90 inhibitor treatment, beginning with anti-nociceptive tolerance. Over a 7-day repeated treatment period, the anti-nociceptive efficacy of morphine steadily decreased in Vehicle treated mice, resulting in a complete loss of anti-nociceptive efficacy by day 7 for the entire 1-10 mg/kg dose range (**Figure 4A-C**). In contrast, KU-32 treatment caused a significant decrease in tolerance over the full 7 day treatment period and full dose range, so that at least some significant anti-nociceptive efficacy remained by day 7 at each dose with KU-32 treatment (**Figure 4A-C**). Notably, baseline responses were not altered by any treatment, demonstrating that KU-32 treatment is not altering baseline nociception (**Figure 4A-C**). To quantify this result, we compared day 1 vs. day 4 responses for each dose and treatment; day 4 was chosen because days 5-7 have no quantifiable Vehicle response for at least one dose each. Dose/response analysis of these data revealed a 21 fold tolerance shift from day 1 to day 4 for Vehicle-treated mice; this tolerance shift was reduced to 2.9 fold in KU-32 treated mice (**Figure 4D**).

**Figure 4:**
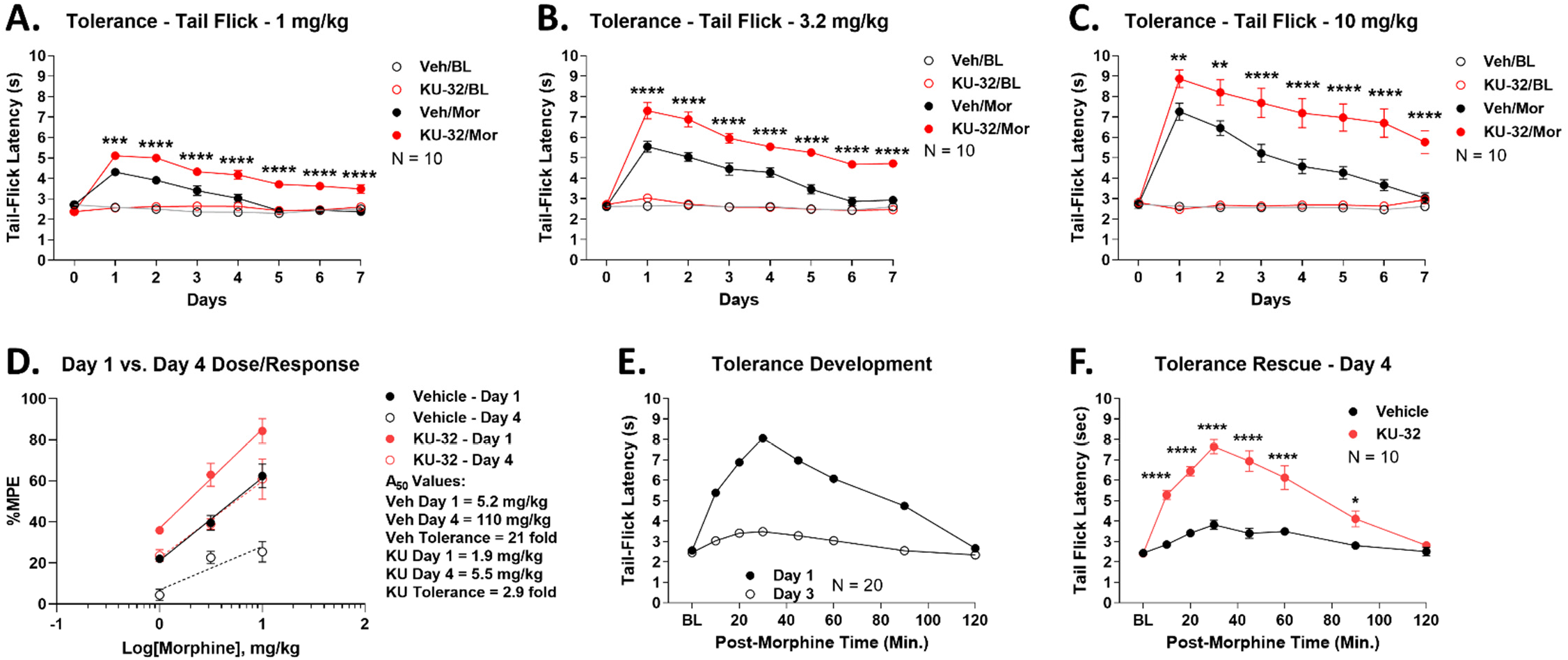
Spinal Hsp90 inhibition reduces morphine anti-nociceptive tolerance and rescues established tolerance. Data reported as the mean ± SEM with sample sizes of mice/group noted in the graphs. **A-C)** Male and female CD-1 mice injected with 0.01 nmol KU-32 or Vehicle control i.t. beginning on day 0 and continuing daily until day 6. The mice were injected with 1-10 mg/kg morphine s.c. twice daily beginning on day 1 and continuing daily until day 7. Thermal tail flick latencies (52°C, 10 sec cutoff) were recorded before treatment on day 0, daily before morphine injection (“BL”), and 30 minutes after each morning morphine injection on days 1-7. **, ***, **** = p < 0.01, 0.001, 0.0001 vs. same time point Veh/Mor group by RM 2 Way ANOVA with Tukey’s *post hoc* test. Individual dose/time curves shown as labeled. KU-32 treatment caused consistent elevation over Vehicle treatment over the 7 day period in all doses. Data not normalized. **D)** Day 1 vs. Day 4 dose/response analysis performed with the data from **A-C**. Data normalized to %MPE (baseline vs. 10 sec cutoff). A_50_: Vehicle Day 1 = 5.2 (3.9 – 7.5) mg/kg, Vehicle Day 4 = 110 (30 – 6300) mg/kg, 21 fold shift; KU-32 Day 1 = 1.9 (1.3 – 2.5) mg/kg, KU-32 Day 4 = 5.5 (3.5 – 12) mg/kg, 2.9 fold shift. **E)** Male and female mice treated with twice daily morphine (10 mg/kg, s.c.) for 3 days to establish tolerance in all mice. Tail flick responses shown on day 1 (acute morphine) and day 3 (morphine-tolerant). **F)** On day 3, after measuring the tail flick time course, the mice were injected with 0.01 nmol KU-32 or Vehicle control, i.t., with a 24 hour recovery. On day 4, the mice were injected again with 10 mg/kg morphine, s.c., and a tail-flick time course performed. Experiment performed in 2 technical replicates. *, **** = p < 0.05, 0.0001 vs. same time point Vehicle group by RM 2 Way ANOVA with Sidak’s *post hoc* test. The KU-32 treated mice, previously tolerant on day 3, showed a statistically significant increase in morphine anti-nociception when compared to Vehicle.

This result suggests that morphine tolerance is blocked by KU-32 treatment; however, KU-32 treatment began before the tolerance regimen and continued during every morphine injection. We thus sought to determine if KU-32 treatment would reverse already-established tolerance. We thus subjected naïve mice to a tolerance regimen as above for 3 days; all mice showed normal morphine anti-nociception on day 1 and a near-complete tolerance by day 3 (**Figure 4E**). We then injected these mice with KU-32 or Vehicle, and 24 hours later, injected morphine again. The Vehicle-treated mice showed little anti-nociception, showing how their tolerance was maintained (**Figure 4F**). In sharp contrast, the KU-32 treated mice showed a full anti-nociceptive response to morphine, comparable to their day 1 response (**Figure 4F**). These results suggest that not only can spinal Hsp90 inhibition reduce the development of tolerance, it can restore responsiveness to already-tolerant mice.

### Spinal Hsp90 inhibition does not alter morphine-induced constipation and reward

Based on our rationale described above, we continued to investigate clinically-relevant opioid side effects. Opioid-induced constipation is a highly clinically significant side effect, that lowers medication compliance and patient quality of life [18]. After KU-32 or Vehicle treatment, we injected a dose-range of morphine and measured fecal output over a 6 hour time course. Compared to saline injected controls, morphine caused ~40% constipation at 1 mg/kg, that plateaued at ~70% constipation at 3.2-10 mg/kg (**Figure 5A-C**). Vehicle vs. KU-32 saline or morphine treatment curves were not significantly different at any dose or time point. After normalizing each treatment group to saline-injected controls, dose/response analysis revealed overlapping curves with a constipation potency of 0.67 (0.32 – 0.94) mg/kg for Vehicle and 0.97 (0.59 – 1.3) mg/kg for KU-32, further supporting the conclusion that KU-32 treatment did not alter morphine constipation (**Figure 5D**).

**Figure 5:**
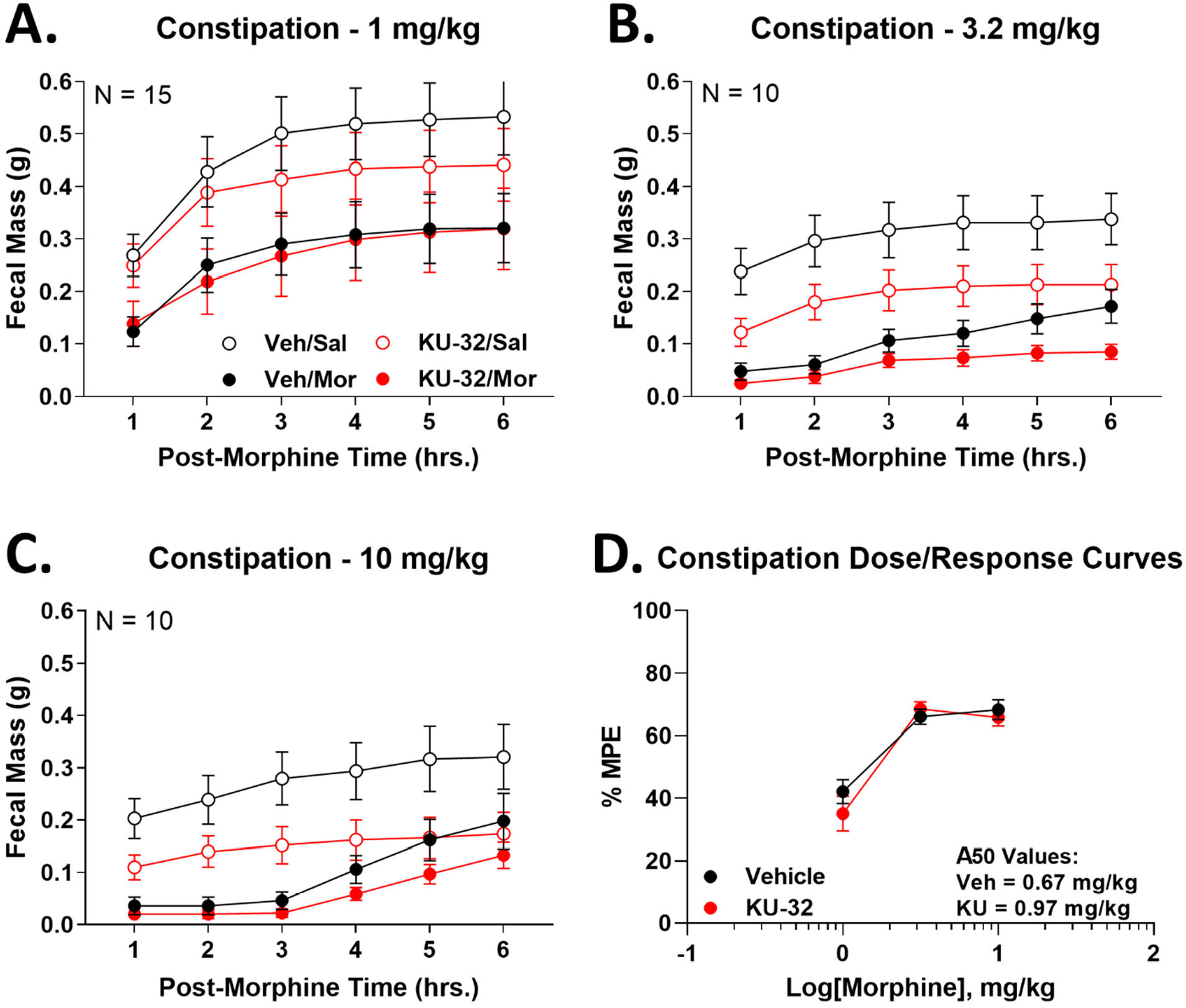
Spinal Hsp90 inhibition does not alter morphine-induced constipation. Male and female CD-1 mice treated with 0.01 nmol KU-32 or Vehicle control i.t., 24 hours, followed by 1-10 mg/kg morphine, s.c.. Fecal mass was measured over 6 hours post-morphine, and used to construct cumulative plots. Curves for morphine as well as saline-injected controls shown. Data reported as the mean ± SEM with the sample size of mice/group noted in the graphs. Experiments performed in 2-3 technical replicates. **A-C)** Individual dose curves reported, along with saline-injected controls. KU-32 treated saline or morphine groups were not statistically different from Vehicle saline or morphine injected groups, respectively, at any time point (p > 0.05). **D)** Morphine injected animals were normalized to saline injected controls for each treatment group (Vehicle or KU-32), and used to construct dose/response curves. The 6 hour area under the curve (AUC) data from **A-C** was further normalized to %MPE (100% MPE = 0% fecal production/100% constipation). Only the 1 and 3.2 mg/kg dose curves were used to calculate the A_50_ since the dose response plateaus between 3.2-10 mg/kg. A_50_: Vehicle = 0.67 (0.32 – 0.94) mg/kg; KU-32 = 0.97 (0.59 – 1.3) mg/kg.

We next tested opioid-induced reward, which is the basis for opioid addiction, and has contributed to an opioid abuse and overdose crisis [17, 39]. We used the well-established conditioned place preference (CPP) assay, which demonstrates reward (or aversion) learning [26]. Over the 3.2-10 mg/kg morphine dose range, we observed an increasing preference for the morphine-paired chamber by Vehicle-treated mice, that rose to the level of significance at 10 mg/kg of morphine (**Figure 6A-C**). KU-32 treatment did not differ significantly from Vehicle treatment at any dose, and also showed significant preference at 10 mg/kg of morphine (**Figure 6A-C**). Dose/response analysis showed overlapping curves and a potency of 5.1 (2.7 – 7.7) mg/kg for Vehicle and 5.3 (-∞ - +∞) mg/kg for KU-32, further supporting a lack of effect of KU-32 treatment on morphine-induced reward learning (**Figure 7D**).

**Figure 6:**
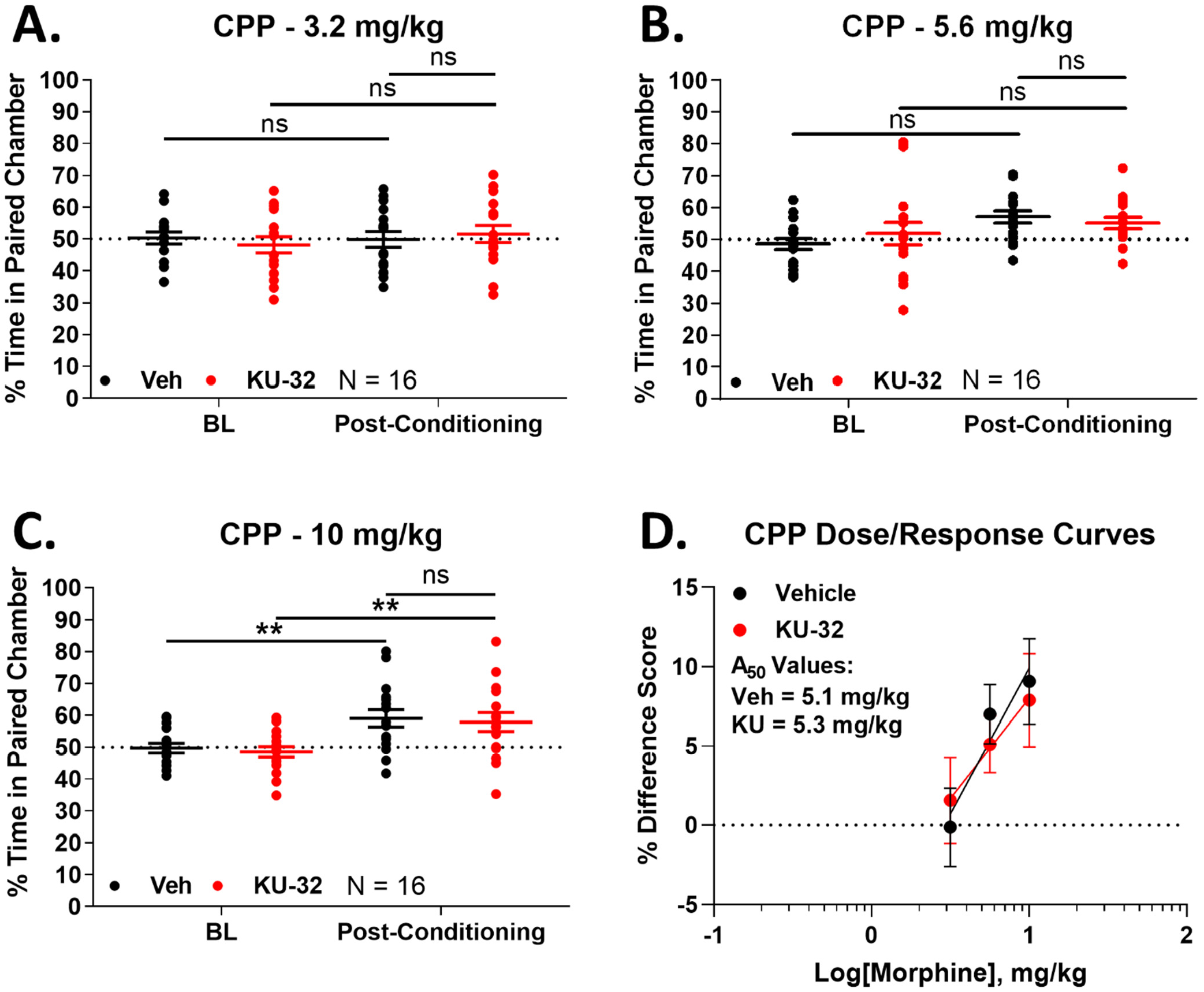
Spinal Hsp90 inhibition does not alter reward learning. Male and female mice were treated over 4 days with KU-32 or Vehicle treatment i.t., combined with 3.2-10 mg/kg morphine, s.c. as a conditioned place preference (CPP) stimulus (see Methods). On day 5, paired chamber preference was recorded. Data reported as the mean ± SEM with the sample size of mice/group noted in the graphs. Each dose performed in 2 technical replicates. ** = p < 0.01 vs. same treatment Baseline (BL) by RM 2 Way ANOVA with Tukey’s *post hoc* test. **AC)** The individual doses are shown. There was a trend to increasing preference with increasing dose that reached significance over baseline at 10 mg/kg for both Vehicle and KU-32 treatment. There was no difference between Vehicle and KU-32 treatment (p > 0.05). **D)** Dose/response analysis was performed, with the data reported as the % Difference Score (see Methods). For the purposes of A_50_ calculation, the response at 10 mg/kg in Vehicle-treated mice was considered to be a maximal response, since 100% preference is not possible in this assay. A_50_: Vehicle = 5.1 (2.7 – 7.7) mg/kg; KU-32 = 5.3 (-∞ - +∞) mg/kg.

**Figure 7:**
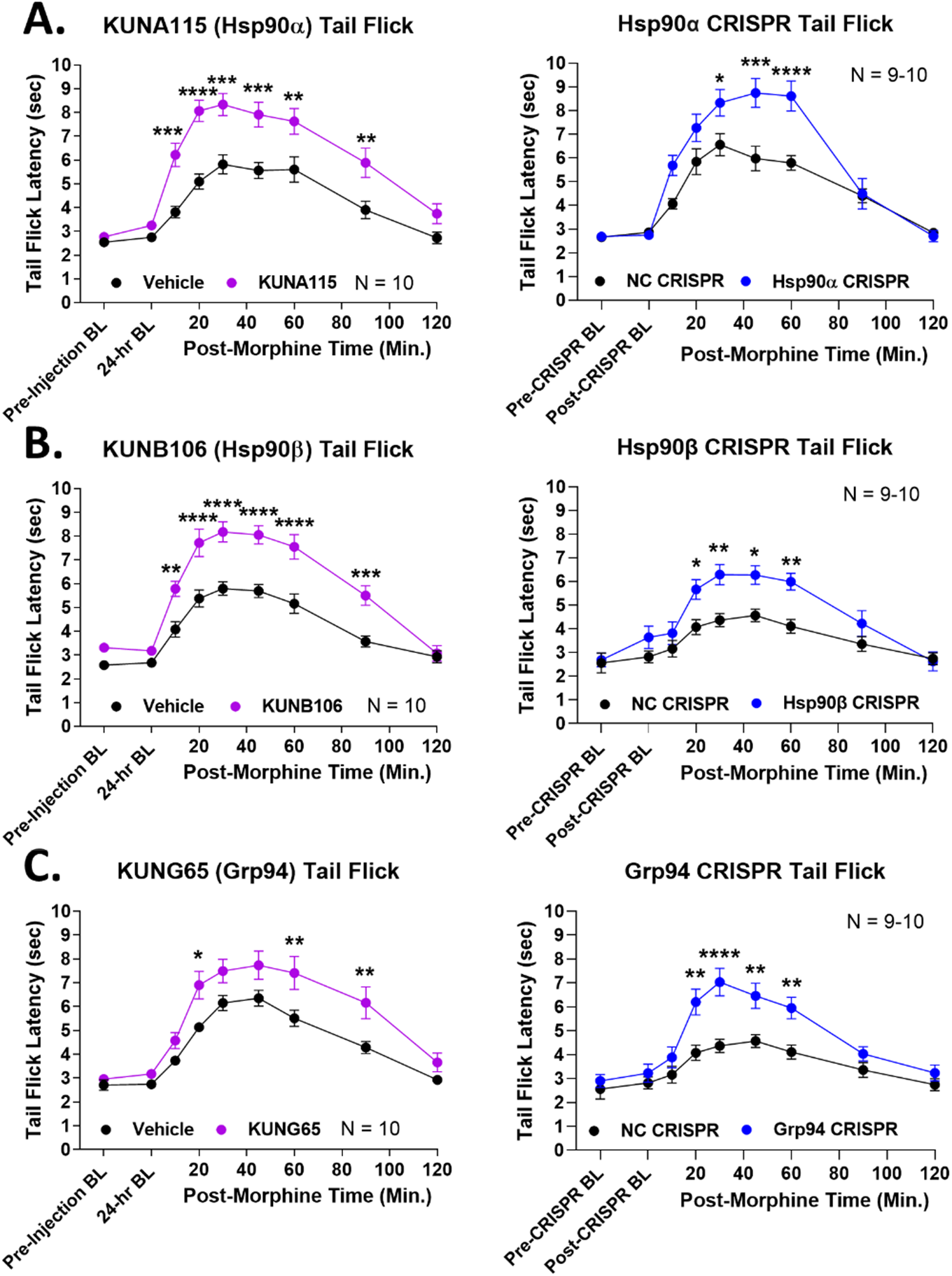
Identification of spinal-cord specific Hsp90 isoforms that regulate opioid anti-nociception. Male and female CD-1 mice were treated with 0.01 nmol of isoform-selective inhibitor or Vehicle control i.t. with a 24 hr treatment time; or with an isoform-selective CRISPR knockdown construct or universal negative control CRISPR construct (NC) with a 10 day treatment time (see Methods). Pre- and post-treatment baselines were recorded using the 52°C tail flick assay (as above), as well as tail flick timecourses in response to 3.2 mg/kg morphine s.c.. Data are presented as the mean ± SEM with the sample size of mice/group noted in each graph. Each experiment was completed in 2 technical replicates. *, **, ***, **** = p < 0.05, 0.01, 0.001, 0.0001 vs. same time point Vehicle/NC group by 2 Way RM ANOVA with Sidak’s *post hoc* test. **A)** The Hsp90α isoform targeted with the selective inhibitor KUNA115 as well as selective CRISPR. **B)** The Hsp90β isoform targeted with the selective inhibitor KUNB106 as well as selective CRISPR. **C)** The Grp94 isoform targeted with the selective inhibitor KUNG65 as well as selective CRISPR. Both methods caused significant anti-nociceptive elevation for all 3 isoforms, confirming that all 3 isoforms regulate opioid anti-nociception in the spinal cord.

### Identification of spinal cord-specific Hsp90 isoforms

The results above confirm our basic hypothesis that inhibition of Hsp90 in the spinal cord can improve the therapeutic index of opioids. However, all experiments above were performed with i.t. injection, which is of limited therapeutic relevance, and we already know that systemic delivery of non-selective Hsp90 inhibitors blocks opioid anti-nociception by inhibiting active Hsp90 in the brain [16]. We thus sought a way to block spinal Hsp90 in a clinically-relevant way. In our earlier work, we found that of the 4 Hsp90 isoforms, only Hsp90α was active in regulating opioid signaling in the brain [14]. We thus hypothesized that if different Hsp90 isoforms were active in the spinal cord, these could be targeted by systemically delivered *selective* inhibitors to selectively block spinal cord Hsp90.

We thus treated mice with selective small molecule inhibitors and targeted CRISPR constructs for each Hsp90 isoform, all delivered by the i.t. route. We found that the Hsp90α-selective inhibitor KUNA115 and Hsp90α-targeted CRISPR enhanced morphine anti-nociception in the tail flick assay (**Figure 7A**). This result suggests that Hsp90α regulates opioid signaling in the brain *and* the spinal cord. However, unlike the brain, inhibitors and CRISPR targeted to Hsp90β and Grp94 also enhanced morphine pain relief (**Figure 7B-C**). This suggests that all 3 Hsp90 isoforms regulate opioid signaling in the spinal cord, while only Hsp90α is active in the brain. If our hypothesis is correct, then systemic inhibition of Hsp90β and Grp94 should recapitulate the benefits of Hsp90 inhibition in the spinal cord by the i.t. route.

### Systemic Grp94 inhibition recapitulates the benefits of spinal Hsp90 inhibition

To test this hypothesis, we used the selective Grp94 inhibitor KUNG65 ([23], **Figure 8A**). KUNG65 is a strongly selective Grp94 inhibitor, with a target K_D_ of 540 nM and at least 73 fold selectivity vs. the other Hsp90 isoforms (**Figure 8B**), making this molecule a good choice for this study. However, we are the first to use KUNG65 *in vivo*, and systemic delivery requires metabolic stability and the ability to cross the blood-brain barrier, which is unknown for this molecule. We thus tested *in vitro* ADME parameters of KUNG65, finding a LogD of 3.3, a solubility of 0.074 μM, and a metabolic half-life of 144 and 32 minutes in human and mouse liver microsomes, respectively (**Figure 8C**). While the solubility is low, the other parameters were promising for *in vivo* delivery. We thus performed a pharmacokinetic study, giving KUNG65 at 10 mg/kg by the oral route, the most challenging route of administration. We found variable but measurable KUNG65 in the plasma, peaking at 79.7 nM in the plasma, and in the spinal cord, peaking at 5.2 nM (**Figure 8D**). While KUNG65 had relatively poor pharmacokinetic performance, preventing the calculation of clearance, half-life, etc., this result was sufficient to demonstrate that the drug could penetrate into the site of action in the spinal cord. We thus delivered KUNG65 at 1 mg/kg by the i.v. route (to enhance spinal cord penetration) in a similar experimental design to the pain assays above, followed by a morphine dose-response in the tail flick assay. Much like our results with KU-32 above, we found that i.v. KUNG65 consistently elevated tail flick anti-nociception in response to morphine across the 1-5.6 mg/kg dose range (**Figure 8E**). We constructed a dose-response curve, finding an A_50_ of 3.6 (2.9 – 4.6) mg/kg in Vehicle treated mice, and 1.9 (1.4 – 2.3) mg/kg in the KUNG65 treated mice, a 1.9 fold shift improvement in morphine potency (**Figure 8F**). This 1.9 fold shift is identical to the fold shift found with i.t. KU-32 treatment in **Figure 1**, providing a strong initial confirmation of our hypothesis. To further evaluate the therapeutic potential of KUNG65, we tested tail flick anti-nociception 48 hours after treatment, instead of 24 hrs as above. This experiment found no effect of KUNG65 treatment, suggesting the effects of the drug persist for 24 but not 48 hrs (**Figure S1**). To rule out confounding effects at the opioid receptors, we also tested for the ability of KUNG65 to bind to any of the human opioid receptors, finding no binding up to a 10 μM concentration (**Figure S2**).

**Figure 8:**
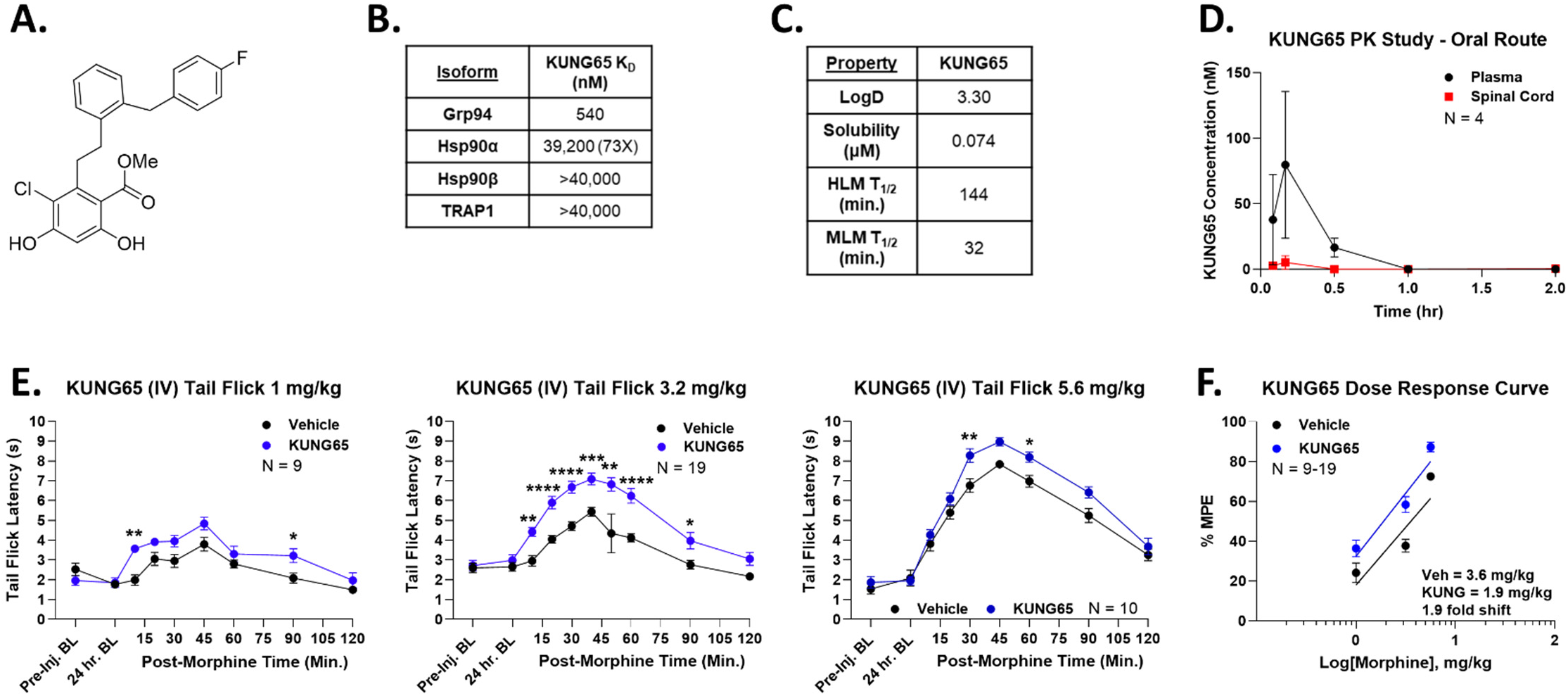
Systemic Grp94 inhibition enhances morphine anti-nociception. **A)** Chemical structure of KUNG65. **B)** The affinity (K_D_) of KUNG65 for each Hsp90 isoform is shown; KUNG65 has a minimum selectivity of 73 fold vs. other isoforms (data taken from [49]). **C)** *In vitro* ADME parameters for KUNG65 are shown (see Methods). HLM/MLM = human/mouse liver microsomes; T_1/2_ = half-life. **D)** A pharmacokinetic (PK) study with KUNG65 was performed, dosed at 10 mg/kg by the oral (p.o.) route in male and female CD-1 mice (see Methods). Data presented as the mean ± SEM, with the sample size of mice/group noted in the graphs, performed in one technical replicate. The KUNG65 is detectable in both plasma and spinal cord, with peak concentrations of 79.7 nM and 5.2 nM, respectively. The drug is undetectable past 30-60 minutes, so PK parameters could not be calculated (half-life, etc.) but the results are sufficient to show that the drug has systemic exposure and blood-brain barrier penetration with oral dosing. **E)** Male and female CD-1 mice treated with KUNG65 (1 mg/kg) or Vehicle injected i.v. with a 24 hr treatment time, followed by 1 – 5.6 mg/kg morphine, s.c., with tail flick time courses performed. Data presented as the mean ± SEM, with the sample size of mice/group noted in the graphs, performed in 2 - 4 technical replicates. *, **, ***, **** = p < 0.05, 0.01, 0.001, 0.0001 vs. same time point Vehicle group by 2 Way RM ANOVA with Sidak’s *post hoc* test. Morphine anti-nociception is consistently elevated, similar to the results with direct spinal inhibition above in **Figure 1**. **F)** The data was transformed into %MPE and used to construct dose/response curves. Vehicle = 3.6 (2.9 – 4.6) mg/kg, KUNG65 = 1.9 (1.4 – 2.3) mg/kg; 1.9 fold improvement in morphine potency.

Continuing our opioid anti-nociception analysis, we then tested i.v. KUNG65 in the post-surgical paw incision model. As expected, i.v. KUNG65 significantly elevated the morphine anti-nociceptive response across the 1 – 3.2 mg/kg dose range (**Figure 9A**). Dose-response analysis showed an A_50_ of 2.0 (1.4 – 3.0) mg/kg in Vehicle treated mice and 0.9 (0.5 – 1.2) mg/kg in the KUNG65 treated mice, a 2.2 fold shift improvement in morphine potency (**Figure 9B**). Again these results support our hypothesis that systemic selective inhibition of Grp94 can mimic the effects of spinal cord Hsp90 inhibition.

**Figure 9:**
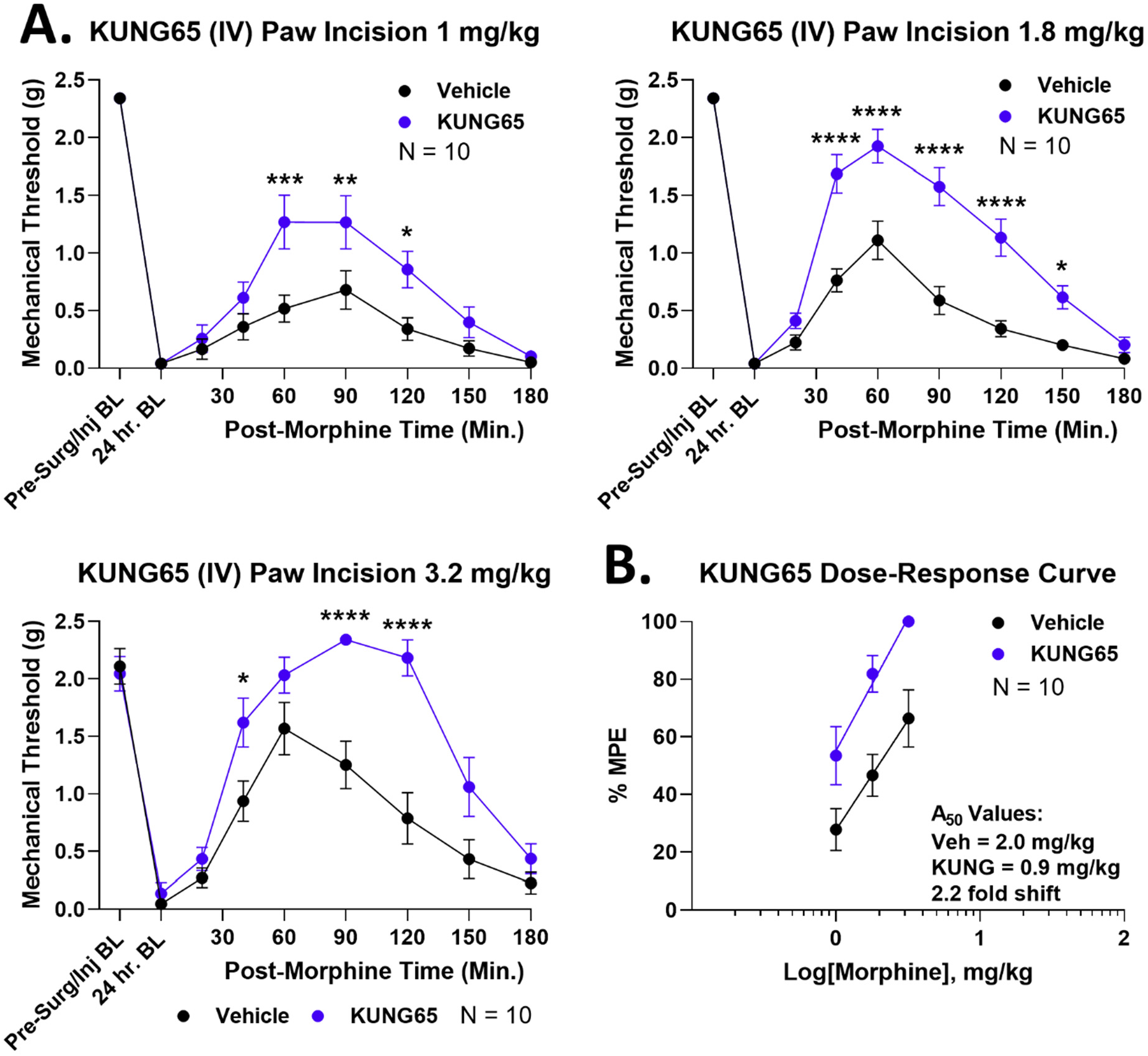
Systemic Grp94 inhibition enhances morphine anti-nociception in paw incision pain. Male and female CD-1 mice treated with paw incision surgery and 1 mg/kg KUNG65 or Vehicle control i.v., 24 hrs, followed by 1 – 3.2 mg/kg s.c. morphine and a Von Frey mechanical allodynia timecourse performed. Data presented as the mean ± SEM, with sample sizes of mice/group noted in each graph; experiments performed in 2 technical replicates per dose. **A)** Individual dose curves shown (no transformation). *, **, ***, **** = p < 0.05, 0.01, 0.001, 0.0001 vs. same time point Vehicle group by 2 Way RM ANOVA with Sidak’s *post hoc* test. KUNG65 treatment consistently elevated anti-nociception. **B)** Data transformed into %MPE and used to construct dose/response curves. A_50_: Vehicle = 2.0 (1.4 – 3.0) mg/kg, KUNG65 = 0.9 (0.5 – 1.2) mg/kg; 2.2 fold improvement in morphine potency.

### Systemic Hsp90β inhibition recapitulates the benefits of spinal Hsp90 inhibition

We continued to test our hypothesis, this time using the Hsp90β-selective inhibitor KUNB106 ([22], **Figure 10A**). Similar to KUNG65 above, KUNB106 is a highly-selective Hsp90β inhibitor, with a target K_D_ of 91 nM and a minimum 275 fold selectivity vs. other isoforms (**Figure 10B**). *In vitro* ADME analysis showed a LogD of 2.26, a solubility of 0.014 μM, and a human and mouse liver microsome half-life of 156 and 63 minutes, respectively (**Figure 10C**). These ADME results and profile are similar to KUNG65 above, so we proceeded directly to *in vivo* testing, using the same 1 mg/kg i.v. dose and route as for KUNG65. Again KUNB106 caused an enhanced morphine tail flick anti-nociceptive response across the entire 1 – 5.6 mg/kg dose range (**Figure 10D**). Upon dose-response analysis, we found an A_50_ of 5.6 (4.4 – ∞) mg/kg for Vehicle treated mice and 1.7 (1.1 – 2.3) mg/kg for KUNB106 treated mice, a 3.3 fold shift (**Figure 10E**). As a test for potential confounds, we tested for the ability of KUNB106 to bind to the opioid receptors, which could explain these results. We did not find any binding to any opioid receptor up to a 10 μM concentration, suggesting our findings are on-target to Hsp90β inhibition (**Figure S2**).

**Figure 10:**
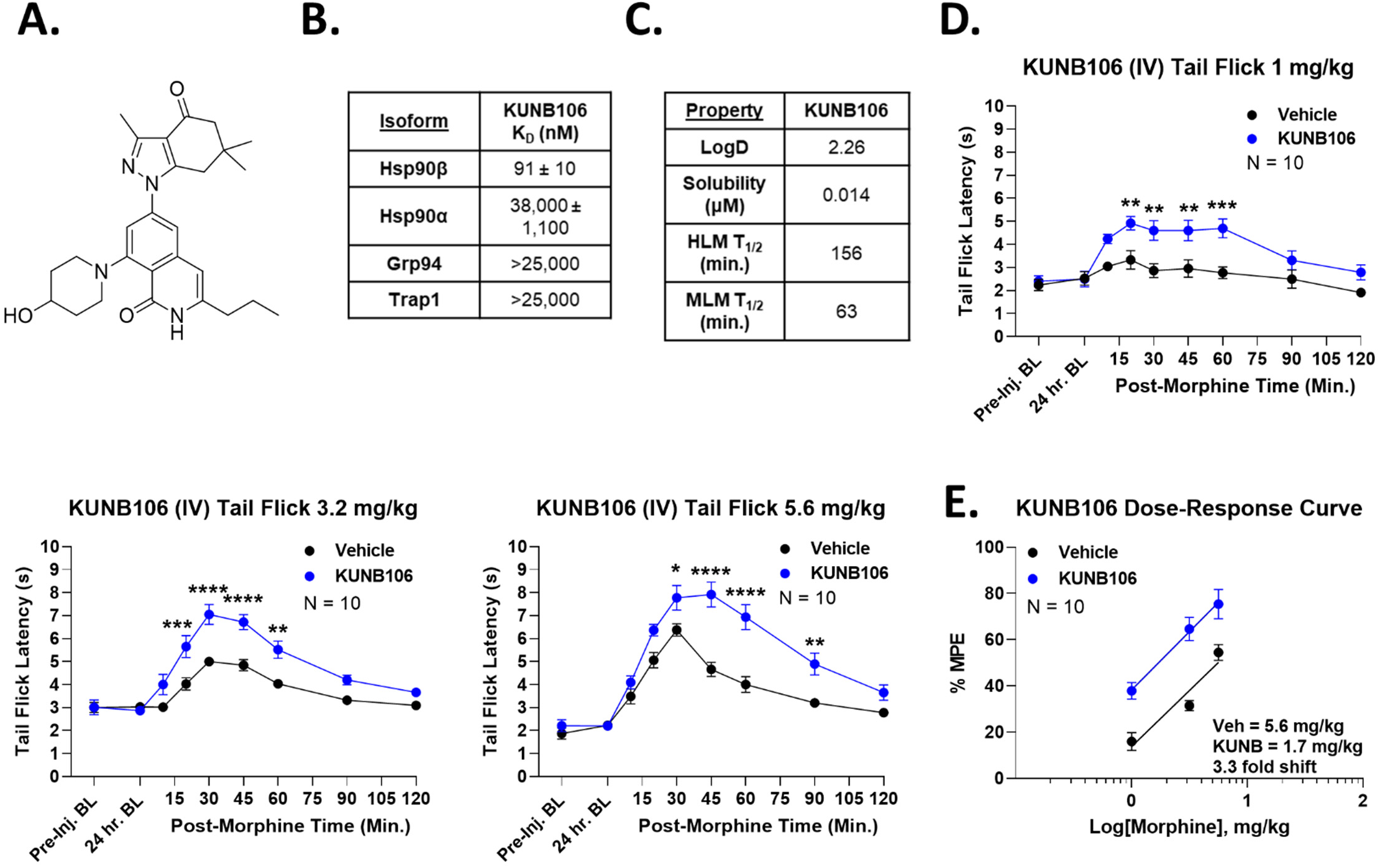
Systemic Hsp90β inhibition enhances morphine anti-nociception. **A)** Chemical structure of the Hsp90β-selective inhibitor KUNB106. **B)** Affinity (K_D_) of KUNB106 for each Hsp90 isoform, showing a minimum selectivity of 275 fold (data taken from [22]). **C)** *In vitro* ADME parameters for KUNB106 are shown (see Methods). HLM/MLM = human/mouse liver microsomes; T_1/2_ = half-life. **D)** Male and female CD-1 mice were treated with 1 mg/kg KUNB106 or Vehicle i.v., 24 hrs, followed by 1 – 5.6 mg/kg morphine s.c. and tail flick timecourses recorded. Data shown as the mean ± SEM, with the sample size of mice per group noted in each graph, completed in 2 technical replicates. *, **, ***, **** = p < 0.05, 0.01, 0.001, 0.0001 vs. same time point Vehicle group by 2 Way RM ANOVA with Sidak’s *post hoc* test. KUNB106 caused increased anti-nociception over the whole morphine dose range. **E)** Dose-response curves generated from the data in **D**, after transformation to %MPE. A_50_: Vehicle = 5.6 (4.4. - ∞) mg/kg, KUNB106 = 1.7 (1.1 – 2.3) mg/kg; 3.3 fold increase in morphine potency.

Extending this analysis again to the post-surgical paw incision model, we found that KUNB106 elevated morphine anti-nociception across the 1 – 3.2 mg/kg dose range (**Figure 11A**). We found an A_50_ of 2.5 (2.0 – ∞) mg/kg for Vehicle treated mice and 0.99 (∞ – ∞) mg/kg for KUNB106 treated mice, a 2.5 fold shift (**Figure 11B**). Together these results support our hypothesis, in that systemic Hsp90β and Grp94 inhibitors both boosted morphine pain relief in a similar manner to direct spinal injection of a non-selective inhibitor like KU-32, suggesting that they are targeting active Hsp90 isoforms in the spinal cord and avoiding active Hsp90 isoforms in the brain.

**Figure 11:**
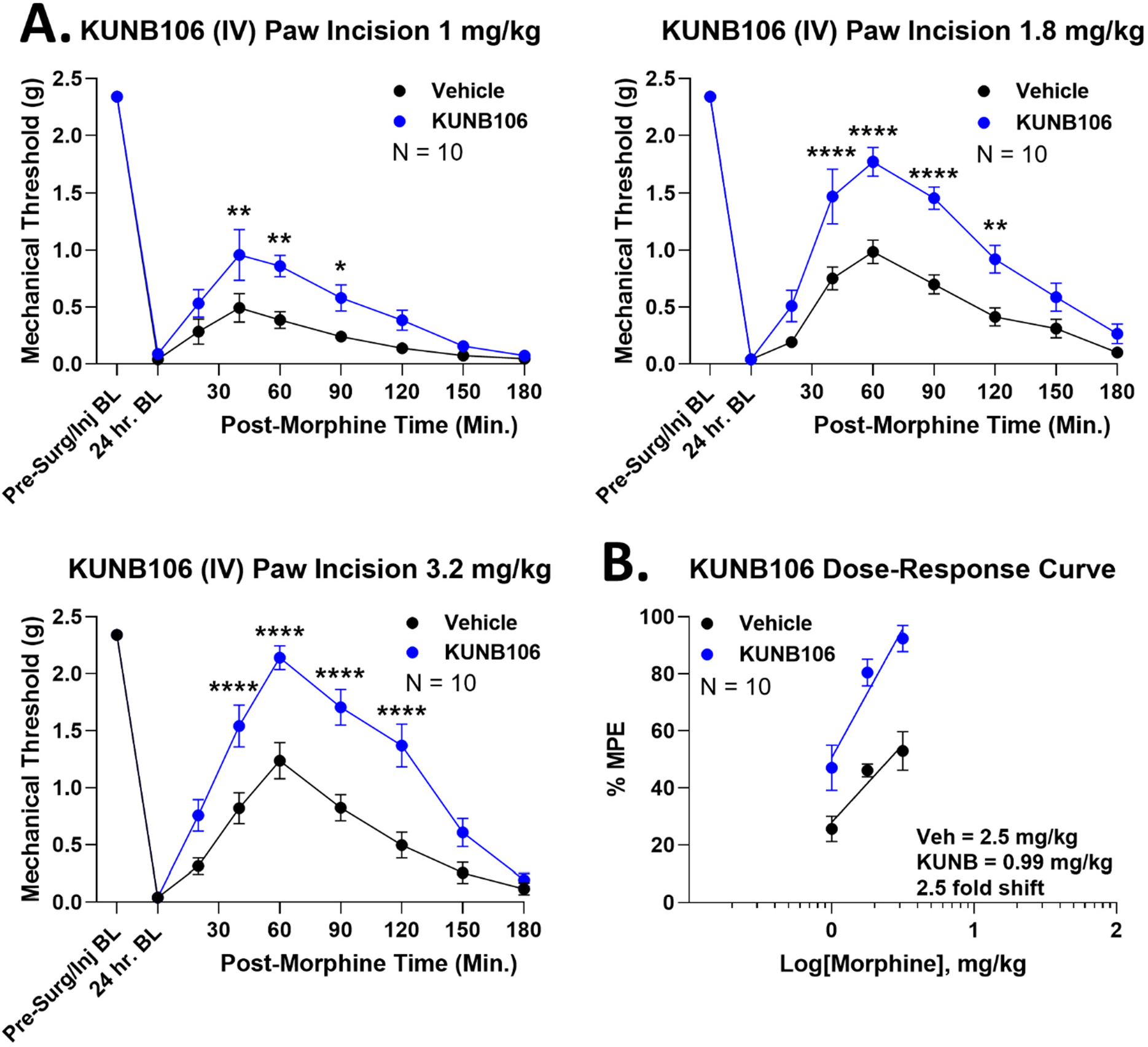
Systemic Hsp90β inhibition enhances morphine anti-nociception in paw incision pain. Male and female CD-1 mice treated with paw incision surgery and 1 mg/kg KUNB106 or Vehicle control i.v., 24 hrs, followed by 1 – 3.2 mg/kg s.c. morphine and a Von Frey mechanical allodynia timecourse performed. Data presented as the mean ± SEM, with sample sizes of mice/group noted in each graph; experiments performed in 2 technical replicates per dose. **A)** Individual dose curves shown (no transformation). *, **, **** = p < 0.05, 0.01, 0.0001 vs. same time point Vehicle group by 2 Way RM ANOVA with Sidak’s *post hoc* test. KUNB106 treatment consistently elevated anti-nociception. **B)** Data transformed into %MPE and used to construct dose/response curves. A_50_: Vehicle = 2.5 (2.0 – ∞) mg/kg, KUNB106 = 0.99 (∞ – ∞) mg/kg; 2.5 fold improvement in morphine potency.

### Systemic Grp94 and Hsp90β inhibitors improve the therapeutic index of morphine

The above results establish that systemic inhibition of Grp94 and Hsp90β boost pain relief much like we saw with direct spinal injection of Hsp90 inhibitor. However, these results do not show that systemic Grp94/Hsp90β inhibitors either improve or do not change side effects, which is necessary to fully show that these treatments boost the therapeutic index of morphine. We thus tested for the impact of KUNG65 and KUNB106 on tolerance and respiratory depression, a key side effect linked to opioid safety and overdose.

We first tested the Grp94 inhibitor KUNG65. We found that acute injection i.v. of 1 mg/kg KUNG65 could rescue established morphine tolerance (**Figure 12A**). This finding was very similar to the tolerance rescue observed with spinal injection of KU-32 in **Figure 4E-F** above, and suggests that KUNG65 can prevent or rescue opioid tolerance. In respiratory depression, we found that KUNG65 had no effect on respiratory activity, either before or after morphine injection (**Figure 12B**). This was an important safety finding, suggesting that while these inhibitors can boost opioid pain relief, they will not boost side effects, thus they improve the therapeutic index of morphine. We found near-identical results for KUNB106, finding that KUNB106 injection rescued established tolerance (**Figure 12C**) while also having no effect on morphine-induced respiratory depression (**Figure 12D**). Together these results confirm that Hsp90β and Grp94 inhibitors do indeed boost the therapeutic index of morphine, enhancing pain relief while either improving or not changing side effects.

**Figure 12:**
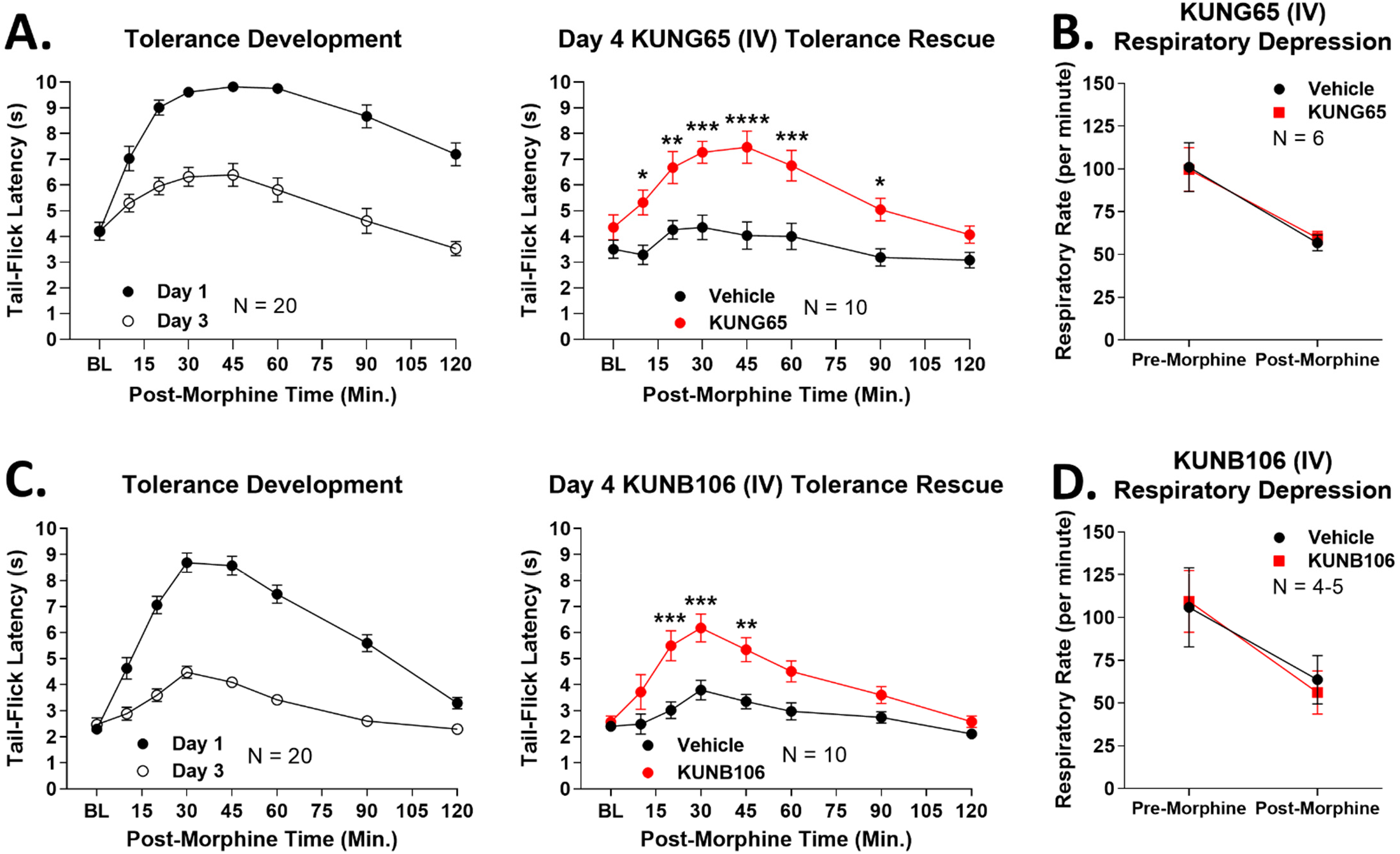
Systemic Grp94 and Hsp90β inhibition rescues established tolerance without worsening opioid-induced respiratory depression. Male and female CD-1 mice used for every experiment, with data presented as the mean ± SEM and the sample size of mice/group noted in each graph. The tolerance experiments were performed in 2 technical replicates, and the respiratory depression experiments in 1 technical replicate. *, **, ***, **** = p < 0.05, 0.01, 0.001, 0.0001 vs. same time point Vehicle group by 2 Way RM ANOVA with Sidak’s *post hoc* test. **A)** Tolerance induced in all mice over 3 days with twice daily injection of 10 mg/kg morphine s.c. (as in **Figure 4**). On day 3, mice injected with 1 mg/kg KUNG65 or Vehicle i.v., 24 hrs, followed by 10 mg/kg morphine s.c. and another tail flick time course. KUNG65 rescued established tolerance much like intrathecal injection of KU-32 above. **B)** Mice injected with 1 mg/kg KUNG65 or Vehicle i.v., 24 hrs, then habituated and baselined in a whole body plethysmography chamber for 30 minutes (see Methods). All mice were then injected with 7.5 mg/kg morphine i.v., and respiratory activity recorded for another hour. KUNG65 had no effect on respiration before or after morphine injection (p > 0.05). **C)** Tolerance induced as above, and on day 3, 1 mg/kg KUNB106 or Vehicle injected i.v., 24 hrs, followed by 10 mg/kg morphine s.c. and another tail flick timecourse. KUNB106 also rescued established tolerance like systemic KUNG65 or intrathecal KU-32. **D)** Mice injected with 1 mg/kg KUNB106 or Vehicle i.v., 24 hrs, then respiratory activity measured as above (including 7.5 mg/kg morphine i.v. challenge). KUNB106 had no impact on respiratory activity before or after morphine (p > 0.05).

### Systemic Hsp90α inhibitor blocks opioid anti-nociception

Finally, we tested the last piece of our hypothesis by systemic i.v. injection of 1 mg/kg KUNA115, a selective Hsp90α inhibitor. If our hypothesis is correct, then this drug should inhibit active Hsp90 in both the brain and spinal cord, leading to a loss of anti-nociception as we saw with systemic non-selective inhibitor. We used this drug in the post-surgical paw incision model, and found that this treatment completely blocked antinociception in response to 3.2 mg/kg morphine (**Figure 13**). This last experiment ties together our model, finding that systemic Hsp90β and Grp94 inhibition recapitulates the beneficial effects of spinal cord inhibition, while Hsp90α inhibition recapitulates the negative effects of brain inhibition.

**Figure 13:**
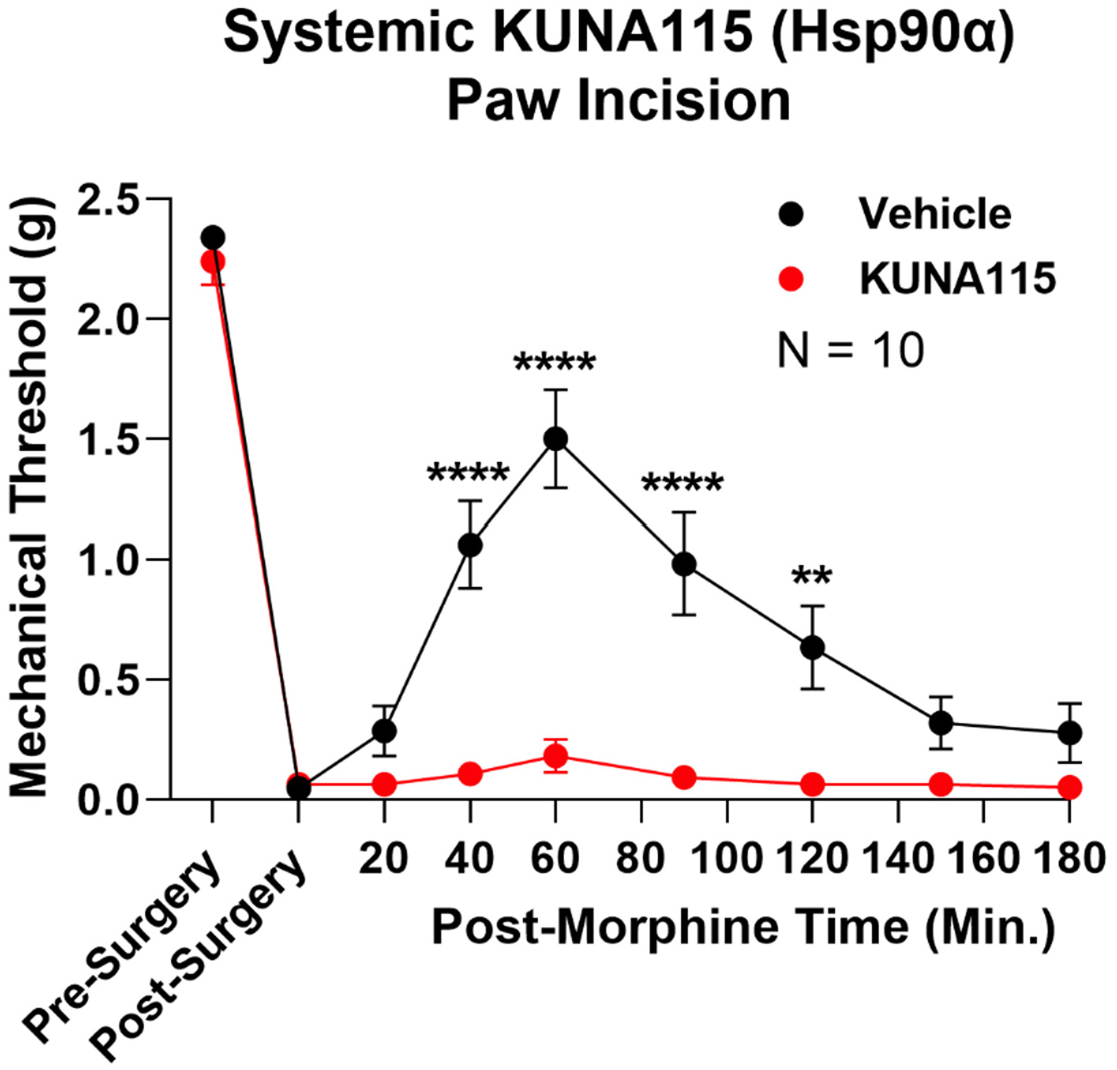
Systemic Hsp90α inhibition blocks opioid anti-nociception. Male and female CD-1 mice had paw incision performed, then injected with 1 mg/kg KUNA115 or Vehicle i.v., 24 hrs, followed by 3.2 mg/kg morphine s.c. and a Von Frey mechanical allodynia timecourse performed. Data presented as the mean ± SEM with the sample size of mice/group noted in the graph (performed in 2 technical replicates). **, **** = p < 0.01, 0.0001 vs. same time point Vehicle group by 2 Way RM ANOVA with Sidak’s *post hoc* test. Unlike Hsp90β and Grp94 inhibition, systemic Hsp90α inhibition via KUNA115 completely blocked opioid anti-nociception in this model.

## Discussion

We show here that Hsp90 inhibition in the spinal cord improves the therapeutic index of morphine, by increasing anti-nociceptive potency by 2-4 fold, reducing and rescuing tolerance, while not changing reward and constipation potency. We have further uncovered a novel strategy to avoid unwanted brain Hsp90 inhibition, which blocks opioid pain relief, by targeting spinal cord-specific Hsp90 isoforms. We found that Hsp90β and Grp94 alone regulate opioid signaling in the spinal cord, while Hsp90α regulates both brain and spinal cord opioid signaling. Thus by delivering isoform-selective Hsp90β and Grp94 inhibitors by a systemic and translationally relevant route, we could enhance opioid pain relief by a similar 2-3 fold while rescuing tolerance and not altering morphine-induced respiratory depression. By contrast, a systemic Hsp90α-selective inhibitor recapitulated brain inhibition, resulting in a loss of opioid anti-nociception. Together these findings establish the potential for Hsp90β and Grp94 inhibitors as opioid co-therapies, which would improve the therapeutic index of opioids, allowing for lower opioid doses with maintained analgesia and reduced side effects. Our model is summarized in **Figure 14**.

**Figure 14:**
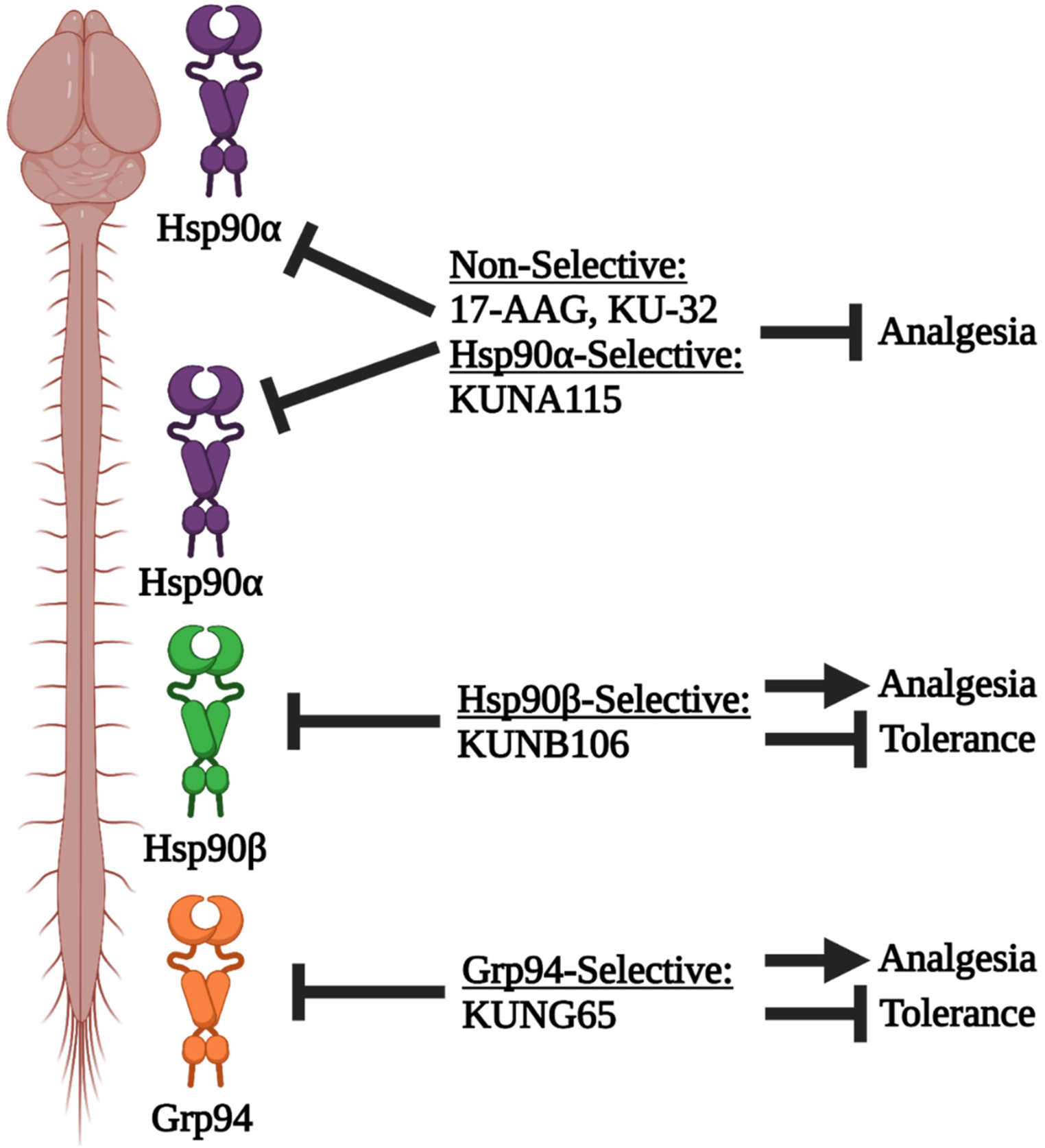
Model of Hsp90 isoform inhibition in the brain vs. spinal cord. In the brain, Hsp90α alone regulates opioid signaling, while in the spinal cord, Hsp90α, Hsp90β, and Grp94 all do so. When Hsp90α is inhibited systemically with non-selective inhibitor or Hsp90α-selective inhibitor, opioid anti-nociception is blocked, since brain Hsp90 is inhibited and a brain-like response is evoked. In contrast, when Hsp90β or Grp94 are inhibited systemically with selective inhibitors, brain inhibition of Hsp90α is avoided, and a spinal cord-like response is evoked – increased anti-nociception and decreased side effects. This model suggests that Hsp90β and Grp94 inhibitors could be given by translationally-relevant routes to improve the therapeutic index of opioids and enable a dose-reduction strategy. Figure created using biorender.com.

Selectively boosting anti-nociceptive potency/efficacy in order to enable a dose-reduction strategy has precedent in the literature. Examples include demonstrated synergy between the cannabinoid receptor type 2 and opioid receptor for anti-nociception, without synergizing the side effects of either [19]. Further examples include selective inhibition of Gβγ signaling downstream of the mu opioid receptor, which similarly potentiates anti-nociception but not side effects [40], morphine and clonidine synergy [41], and using chemokine receptor antagonists to enable opioid dose-reduction [42]. These examples provide precedent and a reason to believe that dose-reduction with Hsp90 inhibitors would work. At the same time, to our knowledge, no such approaches are being used in the clinic, providing novelty and opportunity to use our findings.

Further factors argue for the clinical potential of our findings. Our earlier work suggests that other opioids, specifically oxymorphone, are regulated similarly by Hsp90 inhibition as morphine, albeit in different pain models and inhibitor route [15]. This suggests that a broad spectrum of opioid drugs could be improved. We’ve also shown that these inhibitors have no impact on opioid-induced respiratory depression, which is a key safety concern (**Figure 12**). Our earlier work has shown that non-selective Hsp90 inhibitors given systemically mimic brain inhibition, resulting in a loss of opioid pain relief [16]; this means that our finding that isoform-selective inhibitors can be given systemically to recapitulate spinal inhibition is an important translational advance. Few patients are amenable to an intrathecal delivery route for chronic care, meaning that an isoform-selective approach is necessary to achieve the benefits of spinal Hsp90 inhibition.

One potential concern in adopting these isoform-selective inhibitors for therapy is the potential for on-target side effects. Indeed, early generation non-selective inhibitors like 17-AAG did not make it through clinical trials due to liver toxicity [43]. However, later generation compounds like CNF2024 were far better tolerated, suggesting this toxicity could be off-target rather than Hsp90-mediated [43]. In any case, isoform-selective inhibitors as we propose here should be far better tolerated than non-selective inhibitors, since each isoform has a distinct cellular location and pool of client proteins. This is further supported by a recent report that Hsp90α is the isoform responsible for the serious side effect of retinal degeneration associated with non-selective inhibitors [44]. Happily, our work suggests that Hsp90α is the very isoform that should be avoided for opioid therapy. Together these findings suggest that isoform-selective Hsp90 inhibitors could be feasible for chronic patient therapy.

These observations also raise the question of by what mechanism are these effects taking place. Our earlier work described an ERK-RSK kinase cascade that is not normally active in spinal cord, but becomes “unchained” with Hsp90 inhibition and promotes enhanced anti-nociception [16]. This cascade is presumably responsible for the wider enhancement in anti-nociceptive potency we see here across multiple acute and chronic pain models. At the same time, reward is primarily modulated by a ventral tegmental area-striatal circuit [17] among other forebrain circuits, while constipation is primarily modulated by opioid receptors in the gut [18]; it thus makes sense why local spinal inhibition would not impact these regions and change the potency of those side effects. This leaves tolerance. Several studies have found that spinal circuits modulate opioid tolerance through various mechanisms, suggesting that local spinal inhibition could alter tolerance [45, 46]. Spinal Hsp90 inhibition may favorably alter these mechanisms, blocking tolerance. Similarly, Hsp90 inhibition has been shown to be anti-inflammatory, and spinal neuroinflammation has been shown to contribute to opioid tolerance [7, 47]. Alternately, enhanced anti-nociceptive efficacy could lead to less tolerance over time simply because the efficacy was higher to begin with; in support of this hypothesis, high efficacy opioid agonists have been shown to produce slower/less tolerance than low efficacy agonists [48].

However, these hypotheses only address the slower tolerance seen over time with repeated treatment, they do not explain the tolerance rescue we observed. One potential clue for this rescue mechanism could be our earlier findings that both brain and spinal cord effects of Hsp90 inhibition on anti-nociception require rapid protein translation [14–16]. The translation inhibitor we used had no impact on anti-nociception in Vehicle-treated mice, suggesting that this mechanism is newly activated upon Hsp90 inhibitor treatment. The mechanisms we are uncovering could thus be parallel pathways to the normal/baseline anti-nociceptive signaling, which is further supported by our work on the spinal ERK-RSK cascade [16]. In the case of the tolerance rescue, we could be observing new pathways becoming turned on that have not developed tolerance as have the normal/baseline pathways. These new pathways could thus provide anti-nociceptive responsiveness even when other parts of the system remain tolerant. It’s also unclear at this point whether the tolerance reduction and the tolerance rescue share the same or different mechanisms. Lastly, it is not at all clear why different Hsp90 isoforms are active in brain vs. spinal cord, and whether these isoforms differ in how they regulate opioid signaling. These questions will provide new basic science directions to follow while at the same time these findings can be used to inform a new clinical opioid dose-reduction approach.

## Author Contributions

DID, CSC, KC, and PT performed most mouse experiments, analyzed most of the data, and collaborated on experimental and project design; PT also performed the binding experiments. PB, KAG, and JLB performed some mouse experiments and analyzed some of the data. SM and CB synthesized and purified the novel inhibitors used throughout the paper. DB performed the ADME and pharmacokinetic measurements and analysis. KLH supervised and trained DB in the performance of the ADME/PK assays and provided interpretation of ADME/PK data. BSJB supervised and trained SM and CB in the synthesis of novel inhibitors. JMS conceived the initial idea for the project, collaborated on experimental and project design, analyzed some of the data, and wrote the manuscript. All authors had editorial input into the manuscript.

## Acknowledgments

We would like to acknowledge Drs. Tally Largent-Milnes and Todd Vanderah at the Comprehensive Pain and Addiction Center of the University of Arizona for the use of their conditioned place preference equipment and expertise. This work was supported by an Arizona Biomedical Research Commission New Investigator Award #ADHS18-198875 and institutional funds from the University of Arizona to JMS, as well as R01DA052340 to JMS, KLH, and BSJB. JMS is an equity holder in *Teleport Pharmaceuticals, LLC* and *Botanical Results, LLC*, but these companies are not Hsp90-related. SM and BSJB are equity holders in *Grannus Therapeutic, Inc*. a company focused on the development of Hsp90 inhibitors; the company provided no support or involvement in this study. The authors have no other relevant conflicts of interest to declare.

