## Supplementary Data for "Inhibiting Spinal Cord-Specific Hsp90 Isoforms Reveals a Novel Strategy to Improve the Therapeutic Index of Opioid Treatment"

**Running Title:** Inhibiting Spinal Hsp90 Isoforms Improves Opioid Therapy

**Number of Figures:** 14 main, 2 supplemental

**Figure S1**

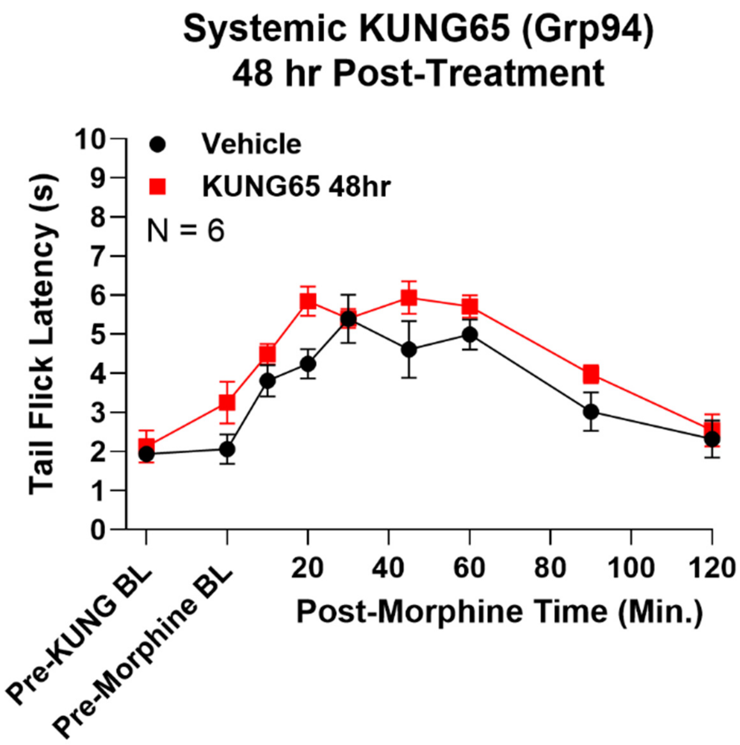


**Figure S1: Enhanced morphine anti-nociception after Grp94 inhibitor treatment dissipates by 48 hours post-treatment.** Male and female CD-1 mice were injected with 1 mg/kg KUNG65 or Vehicle control i.v., followed by 48 hrs of treatment, then 3.2 mg/kg morphine s.c. and a tail flick timecourse. Data displayed as the mean ± SEM with the sample size of mice/group noted in the graph; the experiment was completed with 1 technical replicate. The anti-nociception after 48 hrs of KUNG65 treatment was not significantly different from control ($p > 0.05$), suggesting that the benefits of the treatment wear off between 24 and 48 hrs post-treatment.

**Figure S2**

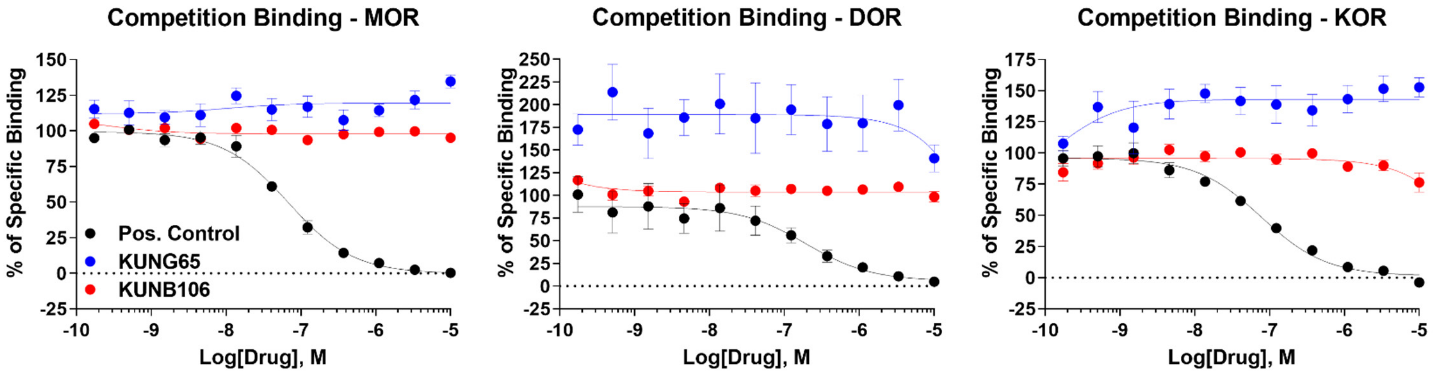


**Figure S2: The Grp94 and Hsp90β inhibitors KUNG65 and KUNB106 do not bind to the opioid receptors.** KUNG65, KUNB106, and positive control (naloxone for mu opioid receptor [MOR] and delta opioid receptor [DOR], U50,488 for kappa opioid receptor [KOR]) were competed against $^{3}$H-diprenorphine at all 3 human opioid receptors using competition radioligand binding (see Methods). Data shown as the mean ± SEM of N = 3 independent experiments. Positive control compounds displayed competition as expected, validating the assay ($K_I$ values: MOR = 34 ± 3 nM; DOR = 120 ± 44 nM; KOR = 37 ± 4 nM). KUNG65 and KUNB106 did not display competition up to a 10 μM concentration, ruling out opioid receptor binding as a potential confound for our findings.
